# Cysteine reactive chloroalkane probe enables HaloTag ligation for downstream chemical proteomics analysis

**DOI:** 10.64898/2026.01.07.698136

**Authors:** Rubaba R. Abanti, Dongqing Wu, Pavel Kielkowski

## Abstract

Chemical proteomics is a powerful method to track proteins labelled by reactive small molecules in living cells on proteome-wide scale. The strategy relies on reactivity and specificity of bioorthogonal ‘click reactions’. Although a variety of bioorthogonal reactions have been developed to facilitate chemical proteomics, their reactivity and specificity might not be comparable with enzymatic reactions. Here we describe an iodoacetamide chloroalkane cysteine reactive probe that is, upon the reaction with nucleophilic cysteine of thioredoxin (TrxA), efficiently and specifically conjugated with HaloTag protein. The TrxA-HaloTag conjugate is utilized for downstream chemoproteomics analysis including in-gel shift assay and mass spectrometry-based proteomics. The TrxA-HaloTag conjugation in whole cell lysate allows fast and efficient pull-down of labelled protein on anti-HaloTag nanobeads resulting in low background after mass spectrometric analysis. The main advantage of the system is its high efficiency and complete biorthogonality due to enzymatic reactivity that is characteristic for HaloTag ligation. The study demonstrates the utility of chloroalkane small compound probes for chemoproteomics applications.

## Introduction

Chemical proteomics is strategy used for deconvolution of non-covalent and covalent interactions between small molecules and proteins on proteome-wide scale. This includes activity-based protein profiling of reactive amino acid residues^1,2^ on proteins using cysteine reactive probes^3,4^ and interrogation of natural products interactions with proteins.^5^ The other application field of chemical proteomics is profiling of protein post-translational modifications that includes protein glycosylation^6– 8^, lipidation^9,10^, ADP-ribosylation^11–13^, AMPylation^14,15^ and tyrosination.^16,17^ The analysis and visualization of the interaction between the chemical proteomic probe and protein is usually done by in-gel fluorescence analysis or by mass spectrometry. However, the critical step of chemical proteomic strategies relies on specificity and reactivity of biorthogonal reactions to conjugate the probe with the reporter or an affinity tag for downstream detection or enrichment of labelled proteins. Therefore, the size and reactivity of the functional group attached to the probe are the main prerequisites for success and specificity of the results in any chemical proteomic study. There are many biorthogonal reactions available including Staudinger ligation^7^, Cu(I)-catalyzed azide-alkyne cycloaddition (CuAAC)^18,19^, strain-promoted azide-alkyne cycloaddition (SPAAC)^20^, inverse electron-demand Diels-Alder (IEDDA) reaction^21^ and azomethine imine-isonitrile reaction^22^ to name a few most common types. They are mainly distinguished by reaction kinetics, size of the functional groups and synthetic accessibility. The chemical methods to establish protein-(probe)-reporter linkage have been complemented by engineered enzymatic systems such as HaloTag^23^, SNAP-tag^24,25^, Connectase^26^ and TUB-tag^27^ as selected examples which all possess fast and specific reactivity characteristic for enzymatic reactions. In particular, HaloTag has been used for diverse applications because of its fast reaction kinetics, stability and relatively small size of the protein, and versatility of the system.^28,29^ Initially, the haloalkane dehalogenase protein DhaA from the bacterial species *Rhodococcus rhodochrous* was engineered in order to result in an irreversible ester bond between a probe and an aspartate’s side chain carboxylic acid of the HaloTag active site.^23^ The HaloTag system was used for many applications including imaging of fusion proteins and affinity protein purification.^28,30^ Recently, HaloTag system was successfully used for chemical proteomic strategy using proteolysis targeting chimeras (PROTACs).^31^ In the PROTAC approach, small compound equipped with a chloroalkane group were designed to target the protein of interest (POI) for proteasomal degradation after conjugation with a HaloTag-E3 ligase fusion protein. Together, these studies underline the robustness and yet untapped application potential of the HaloTag system in chemical proteomics.

In this study, we were inspired by previously developed shift-assay that is based on CuAAC between the alkyne-containing probe labelled protein and azide-polyethylene glycol (PEG) linker in which PEG is branched polymer of five to ten kilodalton in molecular weight resulting in the shift of the labelled protein towards higher mass after separation on sodium dodecyl sulfate-polyacrylamide gel electrophoresis (SDS-PAGE).^32,33^ Here, we designed and synthetized cysteine reactive iodoacetamide chloroalkane probe **1** which was reacted with modified thioredoxin containing only one cysteine. The thioredoxin-chloroalkane probe was characterized by intact protein mass spectrometry and then used for monitoring of conjugation reaction with HaloTag by in-gel shift-assay. The optimized reaction conditions were then tested in cell lysates and pull-down reactions utilizing anti-HaloTag nanobeads followed by LC-MS/MS analysis. Together, our results demonstrate that small compound probes containing chloroalkane tag can be utilized for both gel- and mass spectrometry-based chemical proteomic studies leveraging from high specificity and efficiency of HaloTag ligation.

## Results and Discussion

To initiate the study, the cysteine reactive iodoacetamide chloroalkane probe **1** was synthetized containing linker composed of one ethylene glycol unit and 6-chlorohexyl, which resembles the most used HaloTag ligand TAMRA-chloroalkane (**Figure 1A** and Figure S1).^23^ In order to perform the proof-of-principle experiments showing the utility of HaloTag ligation in chemical proteomics by gel- and mass spectrometry-based analysis (**Figure 1B**), we recombinantly expressed and purified *Escherichia coli* protein TrxA containing a cysteine to serine mutation (TrxA^C33S^) resulting in the protein with a single cysteine residue that can be reacted with the cysteine reactive probe **1** (**Figure 1C** and Supporting Information).^34^ The reaction between TrxA^C33S^ and probe **1** in ratio 1:10 gave about 8-10 % of the modified TrxA^C33S^-chloroalkane as determined by intact protein mass spectrometry (**Figure 1D**). The unmodified TrxA^C33S^ and TrxA^C33S^-chloroalkane were precipitated to remove the excess of the probe **1**. First, concentration-dependent labelling was performed to examine the reactivity of TrxA^C33S^-chloroalkane with the HaloTag. On the Coomassie stained SDS-PAGE, we were delighted to observe the TrxA^C33S^-HaloTag conjugate at about 50 kDa together with HaloTag itself (34 kDa) and TrxA^C33S^ (12 kDa, **Figure 1E**). The successful formation of the TrxA^C33S^-HaloTag conjugate was further verified by Western blot using the anti-TrxA and anti-HaloTag antibodies (**Figure 1E**). Both confirmed the specific formation of the TrxA^C33S^-HaloTag conjugate compared to the negative control experiments, including the free TrxA^C33S^ or HaloTag, only labelled TrxA^C33S^-chloroalkane without HaloTag, non-modified TrxA^C33S^ incubated with HaloTag, and HaloTag treated with probe **1** (**Figure 1E**).

**Figure 1.**
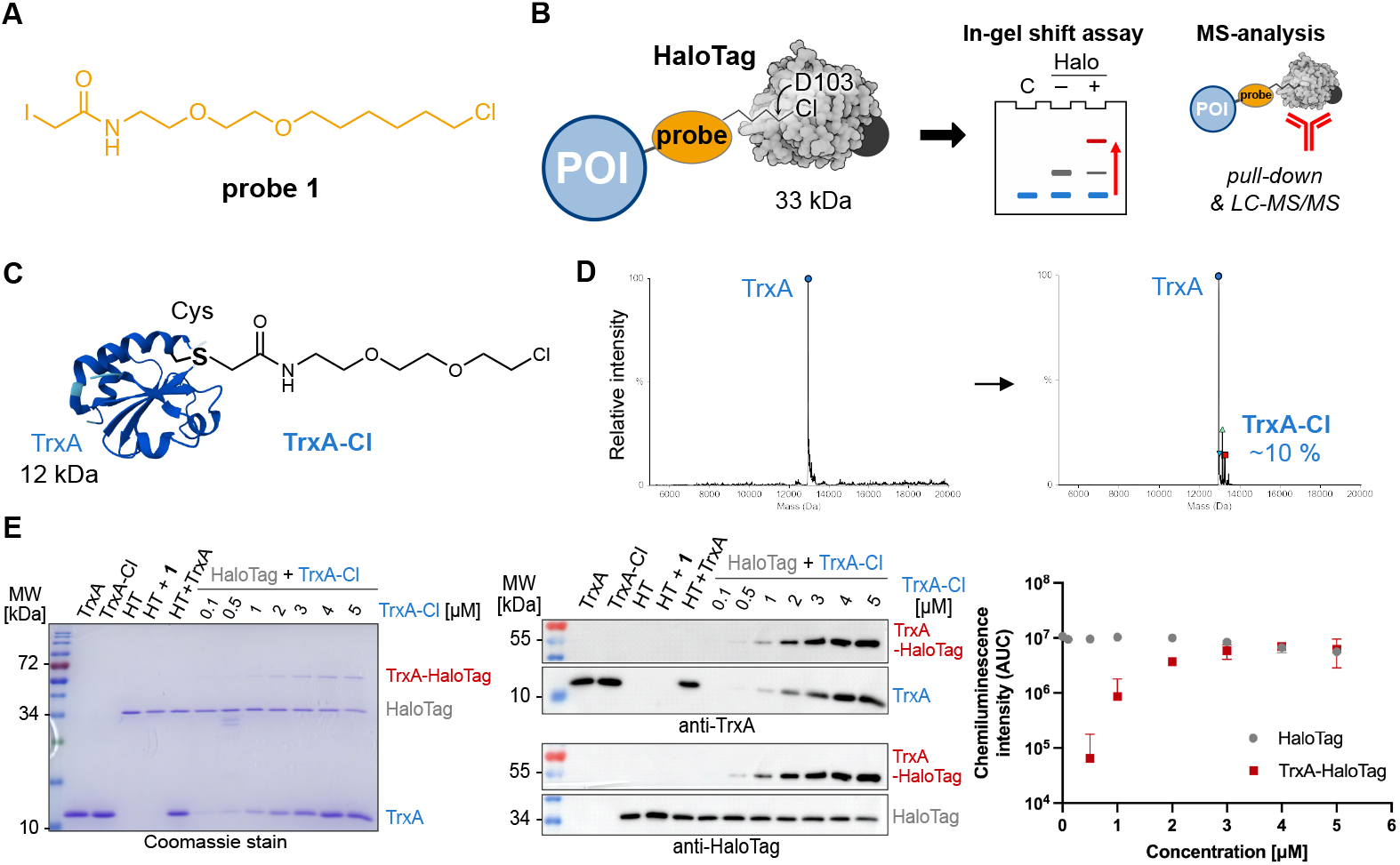
Proof-of-concept: HaloTag application with chemical proteomic probes. **A**) Structure of the cysteine reactive chloroalkane iodoacetamide probe **1. B**) Concept of HaloTag application with chemical proteomic probes for gel- and mass spectrometry-based analysis. **C**) Structure of TrxA^C33S^ labelled with chloroalkane linker. **D**) Intact protein mass spectrometry of TrxA^C33S^ and TrxA^C33S^ modified with the chloroalkane probe **1. E**) Shift-assay of concentration dependent TrxA-chloroalkane conjugation with Halo-Tag (500 nM) analyzed by Coomassie stain (left) and Western blots using anti-TrxA and anti-HaloTag antibodies (middle). The graph (right) shows the quantification of three independent replicates; x-axis is in log_10_ scale; AUC – area under curve. POI – protein of interest. HT – HaloTag.

Next, to further characterize the conjugation reaction conditions, the TrxA^C33S^ / TrxA^C33S^-chloroalkane mixture (5 μM) was titrated with the HaloTag showing the first observable conjugate on Coomassie stain and on Western blots in reaction with 50nM HaloTag (**Figure 2A**). HaloTag reaction with its ligand shows very fast reaction kinetics reaching the diffusion limit of compounds in solution. Therefore, we have tested the time-dependent labelling of TrxA^C33S^-chloroalkane (2 μM) with HaloTag (200 nM, **Figure 2B**). The conjugation product could be observed already after two minutes reaching plateau after ten minutes (**Figure 2B**). After this series of initial experiments with purified proteins, we asked whether the conjugation will be still selective and with the same fast kinetic in complex protein mixtures of cell lysates. To do test that, three different cell lysates from HeLa, HEK293T and SH-SY5Y cells were used as well as varied concentration of TrxA^C33S^-chloroalkane to probe the detection limits. Satisfyingly, in all conditions it was possible to observe specific formation of the TrxA^C33S^-HaloTag conjugate in both Coomassie stained SDS-PAGE and Western blot (**Figure 2C**). In parallel, the time-dependent reaction between TrxA^C33S^-chloroalkane and HaloTag in HEK293T cell lysate confirmed the fast reaction kinetics with clearly observable conjugate formation already after one minute reaction time suggesting even faster reaction kinetics facilitated by surrounding protein-rich environment (**Figure 2D**). To corroborate the anticipated reaction mechanism involving the reactive aspartate of the HaloTag active site, we performed an additional treatment after the TrxA^C33S^-chloroalkane conjugation with HaloTag using the commercial TAMRA-chloroalkane. As expected, we observed decreasing fluorescence signal with increasing TrxA^C33S^-chloroalkane presence in the reaction mixture (**Figure 2E**). Next, although the HaloTag system is compatible with wide range of buffers, we examined the influence of the set of commonly used buffers. The results confirmed the efficient of conjugation in most of the buffers with somewhat decreased reactivity in sodium dodecyl sulfate (SDS) containing buffers (Figure S2, Table S1). Similarly to other chemical biology strategies to conjugate or bring two proteins in close proximity the length of the linker can play a crucial role. The fast reaction kinetics of TAMRA-chloroalkane is dependent on specific interactions between TAMRA-ligand and HaloTag. Probe **1** uses the same linker length as the TAMRA-ligand to fit well into the hydrophobic pocket of the HaloTag to reach the active site and we were wondering, if extending the linker would significantly change the reactivity. Therefore, we have synthetized additional two probes **2** and **3**, each with a prolonged linker length by one ethylene glycol unit (**Figure 2F**). Interestingly, both probes yielded anticipated TrxA^C33S^-HaloTag product in the tested conditions, albeit the buffer seems to have a stronger impact on the overall complex yield in contrast to **1**. Taken together, collectively these experiments demonstrate that HaloTag leads to the productive, specific and fast conjugation with proteins labelled with a reactive small compound probe as exemplified using the cysteine reactive probe **1** allowing to run a gel-based shift-assay to detect the formation of the complex.

**Figure 2.**
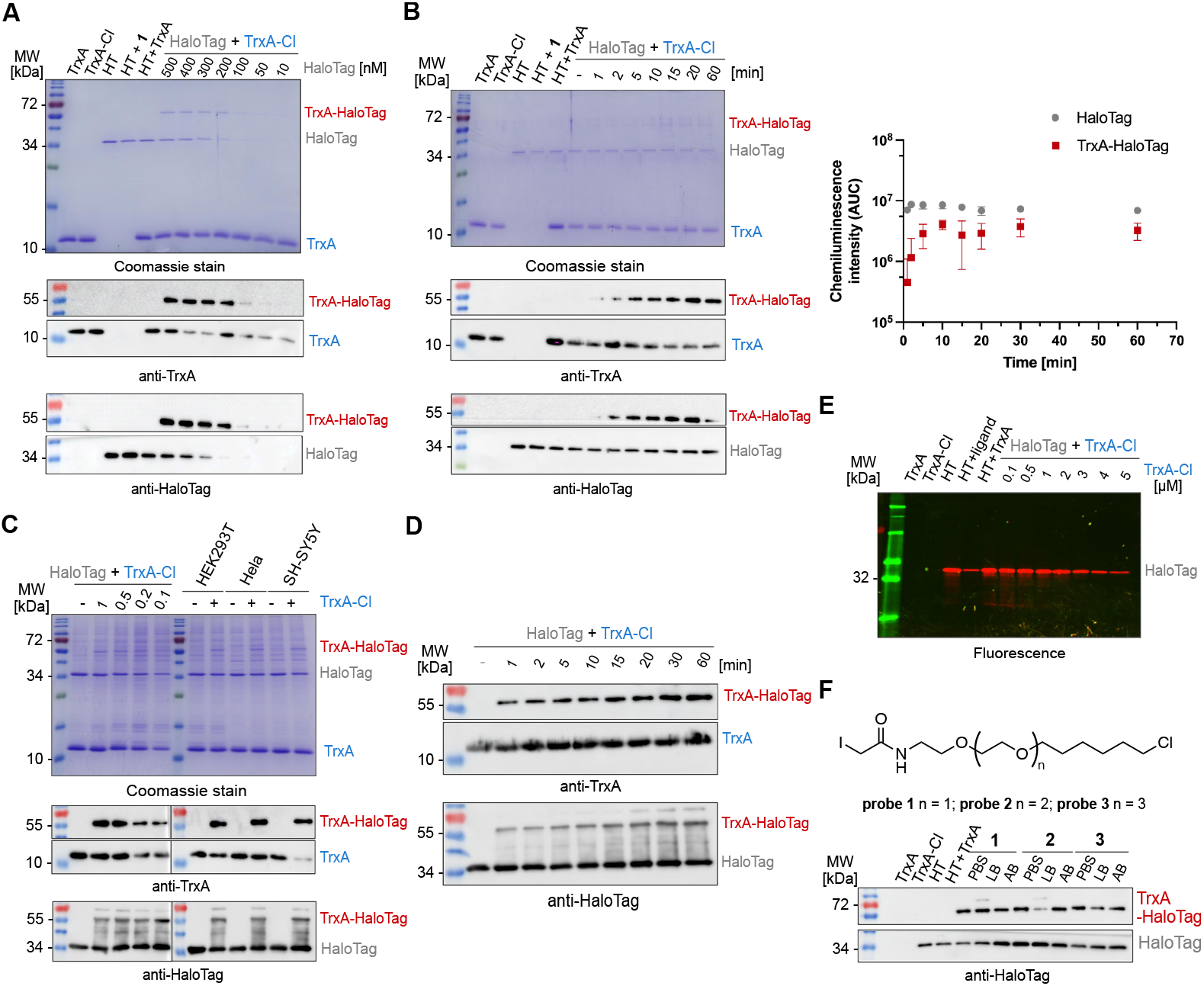
Validation of TrxA^C33S^-chloroalkane conjugation with the HaloTag. **A**) TrxA-chloroalkane (5 μM) conjugation with HaloTag concentration gradient analyzed by Coomassie and Western blot; Reaction conditions: 15 min at 25 °C. **B**) Time-dependent conjugation of TrxA^C33S^-chloralkane (2 μM) with HaloTag (200 nM) analyzed by Coomassie and Western blot (left). The graph (right) shows the quantification of three independent replicates; x-axis is in log_10_ scale; AUC – area under curve. **C**) Conjugation of TrxA^C33S^-chloralkane (5 μM) with HaloTag (500 nM) in cell lysates incubated for 15 min at 25 °C and analyzed by Coomassie stain and Western blot. **D**) Time-dependent conjugation of TrxA^C33S^-chloralkane (5 μM) with HaloTag (500 nM) in cell lysates analyzed by Western blot. **E**) Pulse-chase assay with HaloTag TMR fluorescent ligand: Analysis of HaloTag active site occupancy after the reaction with TrxA^C33S^-chloroalkane using TAMRA-chloroalkane (1 μM) ligand. **F**) Evaluation of the linker length impact on conjugation efficiency with HaloTag (200 nM). Each probe was tested in three different commonly used buffers and analyzed by Western blot. Reaction conditions: TrxA^C33S^-chloroalkane (2 μM) 15 min at 25 °C. PBS – phosphate-buffer saline, LB – lysis buffer, AB – activity buffer. HT – HaloTag.

Typical explorative chemical proteomic experiments involve both gel- and mass spectrometry-based analysis of the probe labelled proteins. To extend the application of HaloTag system in chemical proteomics beyond the gel-based shift-assay, we turned our focus on possibility to pull-down the labelled proteins by utilization of commercially available high-affinity anti-HaloTag magnetic nanobeads. To this end, as described above the TrxA^C33S^-chloroalkane (2 μM) was incubated in HEK293T cell lysate with HaloTag (200 nM) for 15 minutes. After the incubation, the reaction mixture was directly transferred onto the magnetic nanobeads and washed three times with a wash buffer containing Tris-HCl, NaCl and EDTA, with removal of the wash buffer after each step by separation on the magnetic rack (**Figure 3A**). The efficient pull-down of TrxA^C33S^-HaloTag conjugate was confirmed by gel-based analysis (Figure S3). Finally, the enriched protein(s) were proteolytically digested by trypsin and resulting peptides subjected directly to LC-MS/MS analysis. In parallel, control experiments including the lysate incubated either with unmodified TrxA^C33S^ or HaloTag or both were processed in the same way. The analysis of the acquired mass spectra and comparison of all conditions showed highly efficient enrichment of the TrxA^C33S^-HaloTag conjugate with minimal background (**Figure 3B** and Figure S4). Together, this exemplifies the utility of HaloTag conjugates for mass spectrometry-based chemical proteomic pull-down experiments to analyze the small compound probe labelled proteins in complex protein mixtures.

**Figure 3.**
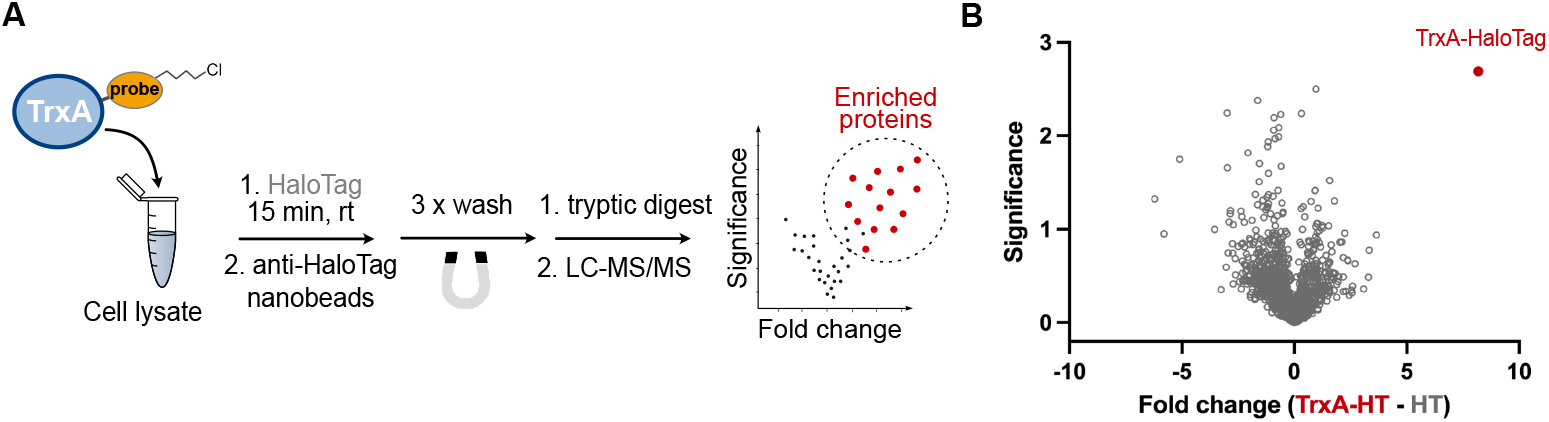
Application of HaloTag for pull-down of labelled protein and mass spectrometry. **A**) General scheme of the workflow. **B**) Volcano plot visualizing pull-down of TrxA^C33S^-HaloTag conjugate from HEK293T cell lysate. Lysate containing HaloTag and TrxA^C33S^ were compared to lysates containing HaloTag and TrxA^C33S^-chloroalkane and both incubated for 15 min at 25 °C. Significance is defined by -log_10_(*p*-value); *p*-value was calculated by Student’s t-test. Fold change is in log_2_ scale. The experiment was done in three independent replicates. HT – HaloTag

In summary, in this study we extend the chemical proteomics toolbox for analysis of proteins labelled with activity-based probes containing the chloroalkane reactive moiety. The strategy leverage from fast and specific reactivity of the HaloTag protein replacing the conventional ‘click reactions’ enabling the downstream analysis by gel-based shift-assay and pull-down of labelled protein followed by mass spectrometry. Principally, the strategy is fully bioorthogonal, fast, specific and can be performed at mild conditions without the necessity for any reaction additives or metal catalysts. Importantly, chloroalkane-based probes and HaloTag are stable and unreactive at virtually all biologically relevant conditions and hence it can be combined with complementary labelling strategies. Based on the recent reports by Shields *et al*. and *Mauker et al*. the chloroalkane linker can be significantly altered beyond the long linear >C10 substrates.^35,36^ We anticipate that both conjugation reaction and pull-down protocol is transferable to other reactive probes and might be combined with many downstream analytical methods including imaging and site-identification experiments that are now unlocked by design and versatility of the HaloTag system.

## Acknowledgements

We are grateful for the support from SFB 1309-325871075 (Deutsche Forschungsgemeinschaft) to P.K. R.A. and P.K. was supported by the BMFTR in the framework of the Cluster4Future program (Cluster for Nucleic Acid Therapeutics Munich, CNATM) (Project ID: 03ZU1201AA). We thank Sabine Schneider for helpful discussions.

