## Supplementary Information for "Cysteine reactive chloroalkane probe enables HaloTag ligation for downstream chemical proteomics analysis"

#### Supplementary Figures

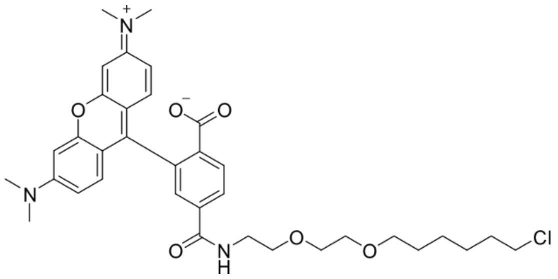

**Figure S1**: Structure of HaloTag-TMR (TAMRA).

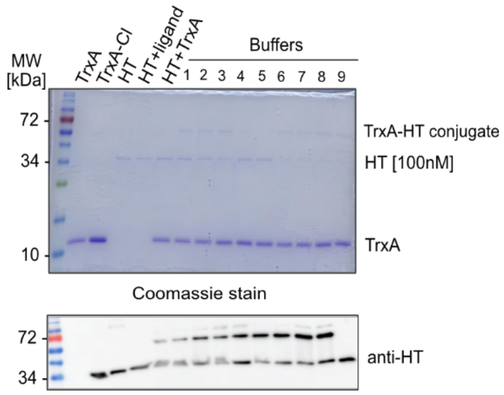

**Figure S2**: TrxA$^{C33S}$-chloroalkane conjugation with HaloTag in different buffer compositions. For 1-9 buffers' composition see Table S1.

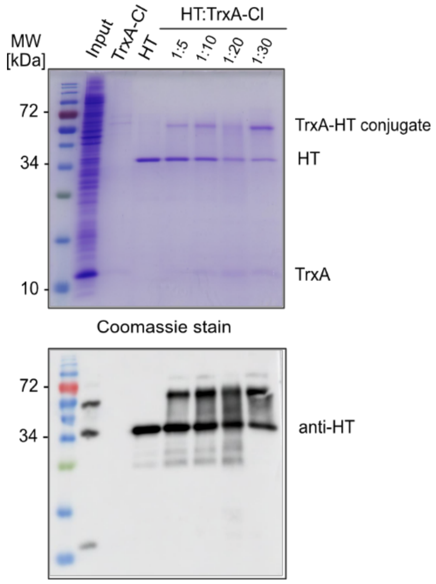

**Figure S3**: In-gel and Western blot analysis of HaloTag and TrxA$^{C33S}$-HaloTag conjugate after pull-down using the Halo-Trap Magnetic Agarose beads.

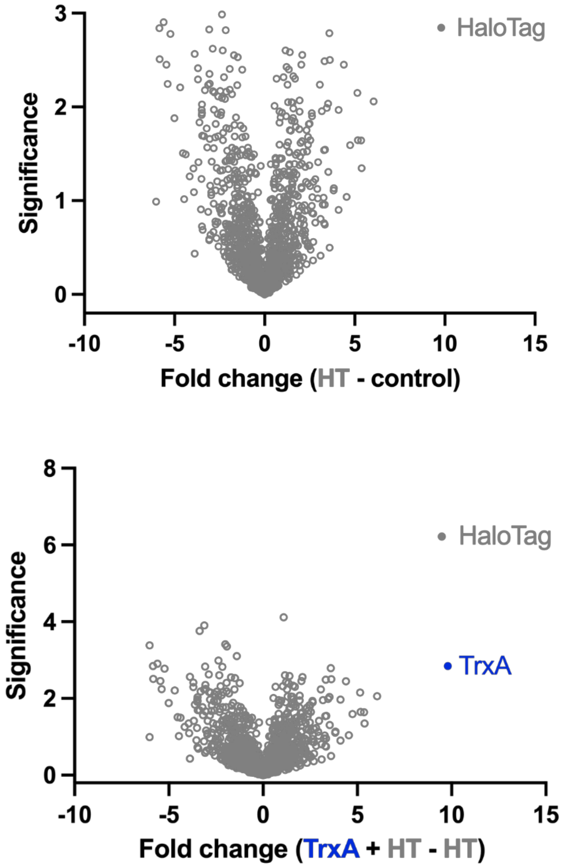

**Figure S4**: LC-MS/MS analysis of HaloTag and modified TrxA$^{C33S}$-HaloTag conjugate after pull-down using the Halo-Trap Magnetic Agarose beads.

#### Supplementary Tables

**Table S1**: Buffer compositions

| | Buffer composition |
|---|---|
| 1 | Phosphate-buffered saline (PBS), pH 7.5 |
| 2 | PBS, 150 mM NaCl, pH 7.4 |
| 3 | PBS, 150 mM NaCl, 0.5 mM EDTA, 0.5% NP-40, pH 7.4 |
| 4 | PBS, 1% NP-40, 0.2% SDS, pH 7.4 |
| 5 | Lysis buffer (20 mM Hepes, 1% NP-40, 0.2% SDS), pH 7.5 |
| 6 | Activity buffer (50 mM Hepes, 150 mM NaCl), pH 7.2 |
| 7 | HNG buffer (50 mM Hepes, 150 mM NaCl, 10% Glycerol), pH 7.3 |
| 8 | 50 mM Hepes, 150 mM NaCl, 0.5mM EDTA, 0.5% NP-40, pH 7.4 |
| 9 | 10 mM Tris/Cl,150 mM NaCl, 0.5mM EDTA, 0.5% NP-40, pH 7.5 |

#### Organic Synthesis

All chemicals, reagents and solvents used for the synthesis were purchased from commercial suppliers and used without further purification.

TLC (thin-layer chromatography) was performed to monitor the reaction progress, which was done on pre-coated silica gel plates (60 F-254, 0.25 mm, from Merck KGaA) as the stationary phase and detected by UV lights (λ = 254 and/or 366 nm).

Flash column chromatography was performed by Pure Chromatography Systems (Pure C-850 FlashPrep) from BÜCHI Labortechnik GmbH on prefilled cartridges (FlashPure EcoFlex Silica) with indicated eluents.

Nuclear magnetic resonance (NMR) spectra ($^1$H and $^{13}$C) were recorded on a Bruker Avance Neo 500 MHz spectrometer. Spectra were analyzed using MestReNova.

MS measurements were obtained from Thermo Scientific ESI-MS based MSQ Plus single quadrupole mass spectrometer.

##### Synthesis of HaloTag ligands

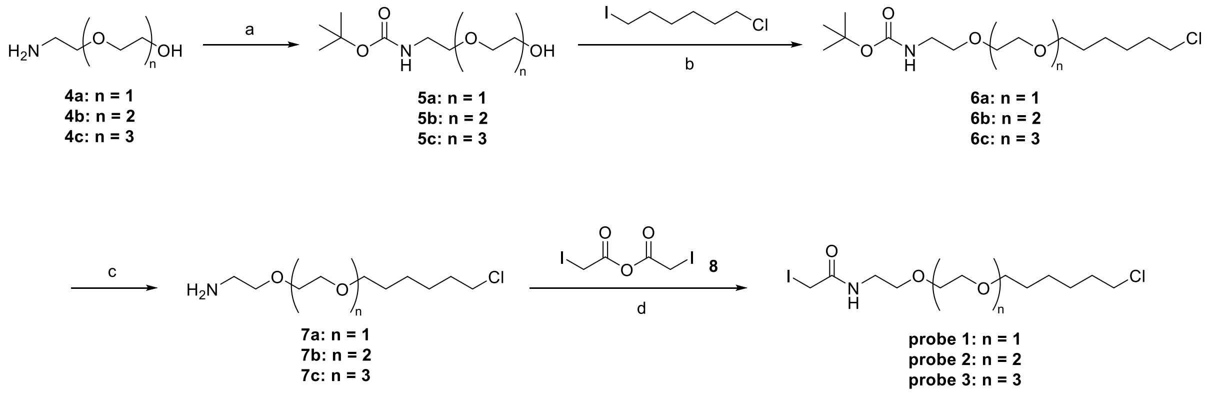

**Scheme S1**: Synthesis of HaloTag ligands (probe 1, 2, 3). Reagents and conditions: (a) $(Boc)_2O$, DCM, rt, overnight; (b) NaH, THF/DMF, 0 °C to rt, overnight; (c) TFA, DCM, 0 °C, 2h; (d) $Et_3N$, DCM, rt, overnight, dark

**General procedure for compound 5:**[1]

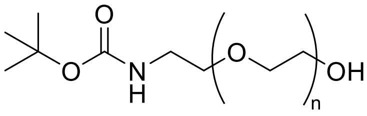

**5a: n = 1**
**5b: n = 2**
**5c: n = 3**

A solution of 0.943 mL of compound **4** (9.51 mmol, 1.00 equiv) in 15 mL of anhydrous DCM was treated with 2.21 g of $(Boc)_2O$ (10.1 mmol, 1.06 equiv) at 0° C and the reaction mixture was stirred for overnight at room temperature. After TLC showed full conversion of the starting material, the solvent was removed under reduced pressure. The resulting product was extracted 3× with 50 mL of DCM from 75 mL of $H_2O$. The combined organic layers were washed with brine and dried over $MgSO_4$, filtered and concentrated *in vacuo*. The crude product was used without further purification in the next synthesis step.

***tert*-butyl (2-(2-hydroxyethoxy)ethyl)carbamate (5a)**

Colorless oil, Yield: 99%.

$^1$H NMR (500 MHz, $CDCl_3$) δ 5.26 (s, 1H), 3.69 – 3.63 (m, 2H), 3.49 (dt, J = 12.4, 5.0 Hz, 4H), 3.26 (q, J = 5.5 Hz, 2H), 3.12 (s, 1H), 1.37 (s, 9H). $^{13}$C NMR (126 MHz, CDCl3) δ 156.20, 79.25, 72.28, 70.27, 61.52, 40.33, 28.38.

MS (ESI, *m/z*): $[M + H]^+$ calculated for $C_9H_{19}NO_4$, 206.13; found: 206.25.

***tert*-butyl (2-(2-(2-hydroxyethoxy)ethoxy)ethyl)carbamate (5b)**

Colorless oil, Yield: 99%.

$^1$H NMR (500 MHz, CDCl3) δ 5.13 (s, 1H), 3.71 – 3.65 (m, 2H), 3.62 – 3.52 (m, 6H), 3.49 (s, 2H), 3.26 (s, 2H), 2.68 (t, J = 6.2 Hz, 1H), 1.38 (s, 9H). $^{13}$C NMR (126 MHz, CDCl3) δ 156.03, 79.26, 72.58, 70.39, 70.28, 70.23, 61.69, 40.32, 28.40.

MS (ESI, *m/z*): $[M + H]^+$ calculated for $C_{11}H_{23}NO_5$, 250.16; found: 250.28.

***tert*-butyl (2-(2-(2-(2-hydroxyethoxy)ethoxy)ethoxy)ethyl)carbamate (5c)**

Colorless oil, Yield: 99%.

$^1$H NMR (500 MHz, CDCl3) δ 5.56 (s, 1H), 3.69 – 3.62 (m, 4H), 3.61 – 3.53 (m, 8H), 3.47 (t, J = 5.0 Hz, 2H), 3.25 (q, J = 5.4 Hz, 2H), 2.97 (t, J = 6.3 Hz, 1H), 1.38 (s, 9H). $^{13}$C NMR (126 MHz, CDCl3) δ 156.21, 78.97, 72.64, 70.64, 70.47, 70.28, 70.10, 61.68, 40.40, 28.46.

MS (ESI, *m/z*): $[M + H]^+$ calculated for $C_{13}H_{27}NO_6$, 294.18; found: 294.32.

**General procedure for compound 6:**[1]

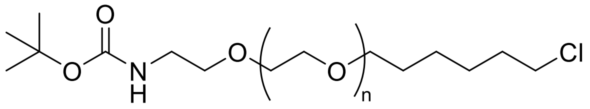

**6a: n = 1**
**6b: n = 2**
**6c: n = 3**

A solution of compound **5** (1.88 g, 9.16 mmol, 1.00 equiv) in 27 mL of THF/DMF 2/1 was treated with sodium hydride 60% in mineral oil (330 mg, 13.7 mmol, 1.50 equiv) portionwise at 0 °C in an ice bath. After stirring for 30 min, 1-chloro-6-iodohexane (3.16 g, 1.99 mL, 12.8 mmol, 1.40 equiv) was added to the reaction mixture at 0 °C and was stirred for 30 min and then at room temperature for 18 h. The reaction mixture was quenched at 0 °C by the addition of aq. sat. $NH_4Cl$. The aqueous phase was extracted 3× with 25 mL of ethyl acetate, and the combined organic layers were washed with brine. The organic layer was dried over $MgSO_4$, filtered and concentrated *in* vacuo. The residue was purified by column chromatography (eluent: hexane/ethyl acetate, 4:1) to obtain compound **6** as colorless oil.

***tert*-butyl (2-(2-((6-chlorohexyl)oxy)ethoxy)ethyl)carbamate (6a)**

Colorless oil, Yield: 33%.

$^1$H NMR (500 MHz, CDCl3) δ 4.92 (s, 1H), 3.49 – 3.46 (m, 2H), 3.45 – 3.38 (m, 6H), 3.33 (t, J = 6.7 Hz, 2H), 3.18 (d, J = 5.8 Hz, 2H), 1.68 – 1.61 (m, 2H), 1.51 – 1.44 (m, 2H), 1.31 (s, 11H), 1.28 – 1.22 (m, 2H). $^{13}$C NMR (126 MHz, CDCl3) δ 155.98, 79.09, 71.24, 70.24, 70.18, 70.01, 44.99, 40.32, 32.51, 29.41, 28.40, 26.66, 25.39.

MS (ESI, *m/z*): $[M + H]^+$ calculated for $C_{15}H_{30}ClNO_4$, 324.18; found: 324.29.

***tert*-butyl (2-(2-(2-((6-chlorohexyl)oxy)ethoxy)ethoxy)ethyl)carbamate (6b)**

Colorless oil, Yield: 34%.

$^1$H NMR (500 MHz, CDCl3) δ 4.98 (s, 1H), 3.60 – 3.50 (m, 8H), 3.46 (t, J = 6.7 Hz, 4H), 3.40 (t, J = 6.7 Hz, 2H), 3.24 (d, J = 5.4 Hz, 2H), 1.74 – 1.67 (m, 2H), 1.57 – 1.49 (m, 2H), 1.38 (s, 11H), 1.34 – 1.27 (m, 2H). $^{13}$C NMR (126 MHz, CDCl3) δ 156.00, 79.11, 71.25, 70.64, 70.55, 70.27, 70.22, 70.11, 45.02, 40.37, 32.54, 29.44, 28.42, 26.69, 25.42.

MS (ESI, *m/z*): $[M + H]^+$ calculated for $C_{17}H_{34}ClNO_5$, 368.21; found: 368.43.

**_tert_-butyl (18-chloro-3,6,9,12-tetraoxaoctadecyl)carbamate (6c)**

Colorless oil, Yield: 33%.

$^1$H NMR (500 MHz, CDCl3) δ 4.99 (s, 1H), 3.62 – 3.53 (m, 10H), 3.52 (d, J = 5.6 Hz, 2H), 3.46 (t, J = 6.8 Hz, 4H), 3.39 (t, J = 6.7 Hz, 2H), 3.24 (d, J = 5.4 Hz, 2H), 1.74 – 1.67 (m, 2H), 1.56 – 1.49 (m, 2H), 1.38 (s, 11H), 1.34 – 1.27 (m, 2H). $^{13}$C NMR (126 MHz, CDCl3) δ 156.01, 79.13, 71.24, 70.63, 70.62, 70.55, 70.27, 70.24, 70.12, 45.04, 40.39, 32.56, 29.46, 28.43, 26.70, 25.43.

MS (ESI, *m/z*): $[M + H]^+$ calculated for $C_{19}H_{38}ClNO_6$, 412.24; found: 412.41.

**General procedure for compound 7:[1]**

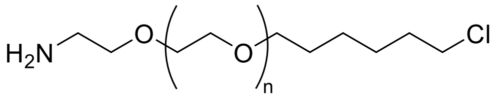

**7a: n = 1**
**7b: n = 2**
**7c: n = 3**

To a solution of compound **6** (807 mg, 2.49 mmol, 1.00 equiv.) in DCM (18 mL) at 0 °C were added TFA (3 mL). After stirring at 0 °C for 2.5 h, solvents were removed under reduced pressure and the residue was diluted with MeOH (18 mL). The solution was cooled to 0 °C and $K_2CO_3$ (988 mg, 7.15 mmol, 2.87 equiv.) was added to the mixture. The mixture was stirred at the same temperature for 10 min, filtered, and evaporated. The residue was diluted with $H_2O$ (12 mL) and the mixture was extracted 4x with ethyl acetate. The combined organic layers were dried over $MgSO_4$, filtered, and concentrated *in vacuo* to obtain compound **7**.

**2-(2-((6-chlorohexyl)oxy)ethoxy)ethan-1-amine (7a)**

Yellow oil, Yield: quant.

$^1H$ NMR (500 MHz, CDCl3) δ 7.84 (s, 3H), 3.70 – 3.63 (m, 2H), 3.62 – 3.55 (m, 2H), 3.53 – 3.48 (m, 2H), 3.45 (t, J = 6.7 Hz, 2H), 3.39 (t, J = 6.9 Hz, 2H), 3.10 (s, 2H), 1.73 – 1.65 (m, 2H), 1.51 (dd, J = 14.9, 7.1 Hz, 2H), 1.41 – 1.33 (m, 2H), 1.30 – 1.23 (m, 2H). $^{13}C$ NMR (126 MHz, CDCl3) δ 71.27, 70.24, 69.72, 66.50, 45.00, 39.71, 32.43, 29.16, 26.58, 25.21.

MS (ESI, *m/z*): $[M + H]^+$ calculated for $C_{10}H_{22}ClNO_2$, 224.13; found: 224.25.

**2-(2-(2-((6-chlorohexyl)oxy)ethoxy)ethoxy)ethan-1-amine (7b)**

Yellow oil, Yield: 92%.

$^1H$ NMR (500 MHz, CDCl3) δ 4.00 (s, 2H), 3.62 – 3.49 (m, 10H), 3.46 (t, J = 6.7 Hz, 2H), 3.39 (t, J = 6.7 Hz, 2H), 2.90 (t, J = 5.1 Hz, 2H), 1.75 – 1.66 (m, 2H), 1.57 – 1.48 (m, 2H), 1.42 – 1.34 (m, 2H), 1.34 – 1.26 (m, 2H). $^{13}C$ NMR (126 MHz, CDCl3) δ 71.24, 70.95, 70.50, 70.46, 70.16, 69.95, 45.06, 40.89, 32.53, 29.37, 26.67, 25.37.

MS (ESI, *m/z*): $[M + H]^+$ calculated for $C_{12}H_{26}ClNO_3$, 268.16; found: 268.31.

**18-chloro-3,6,9,12-tetraoxaoctadecan-1-amine (7c)**

Yellow oil, Yield: 85%.

$^1H$ NMR (500 MHz, CDCl3) δ 3.62 – 3.55 (m, 10H), 3.54 – 3.49 (m, 4H), 3.46 (t, J = 6.7 Hz, 2H), 3.40 (t, J = 6.7 Hz, 2H), 3.12 (s, 2H), 2.85 (t, J = 5.1 Hz, 2H), 1.74 – 1.67 (m, 2H), 1.56 – 1.49 (m, 2H), 1.42 – 1.35 (m, 2H), 1.33 – 1.26 (m, 2H). $^{13}C$ NMR (126 MHz, CDCl3) δ 72.23, 71.27, 70.53, 70.51, 70.48, 70.36, 70.10, 45.04, 41.41, 32.53, 29.42, 26.68, 25.38.

MS (ESI, *m/z*): $[M + H]^+$ calculated for $C_{14}H_{30}ClNO_4$, 312.19; found: 312.34.

**General procedure for probe 1-3:**

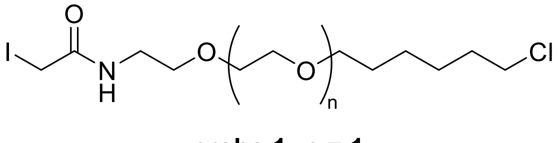

**probe 1: n = 1**
**probe 2: n = 2**
**probe 3: n = 3**

Iodoacetic anhydride (**8**)[2] was synthesized by adding 1.19 g of iodoacetic acid (6.4 mmol) dissolved in 20 mL ethyl acetate to 660 mg of EDC.HCl (3.2 mmol) dissolved in an additional 20 mL of ethyl acetate. The reaction was allowed to stir for 2 hours in the dark. The solution was extracted with water and the organic layers were washed with brine and dried with $MgSO_4$, filtered, and concentrated *in vacuo*.

270 mg of amine (1.2 mmol) was added to 10 mL of acetonitrile with 500 µL triethylamine (3.6 mmol) and stirred for 1 hour to dissolve, resulting in a clear, dark solution. Iodoacetic anhydride (**8**) was then dissolved in 5 mL acetonitrile and added to the amine solution and the mixture was stirred overnight at room temperature in the dark. The solvents were evaporated *in* vacuo and the residue was purified by column chromatography (eluent: ethyl acetate/MeOH, 20:1) to obtain desired probe as yellow oil.

**_N_-(2-(2-((6-chlorohexyl)oxy)ethoxy)ethyl)-2-iodoacetamide (Probe 1)**

Yellow oil, Yield: 11%.

$^{1}$H NMR (500 MHz, CDCl3) δ 3.62 (m, 2H), 3.57 (ddt, J = 5.7, 3.5, 1.7 Hz, 2H), 3.54 – 3.49 (m, 4H), 3.47 (td, J = 6.7, 1.9 Hz, 2H), 3.44 – 3.35 (m, 4H), 1.71 (dtd, J = 14.4, 6.7, 2.3 Hz, 2H), 1.59 – 1.51 (m, 2H), 1.43 – 1.36 (m, 2H), 1.35 – 1.28 (m, 2H). $^{13}$C NMR (126 MHz, CDCl3) δ 167.61, 71.98, 71.06, 70.73, 70.02, 45.70, 40.81, 33.18, 30.15, 27.35, 26.09.

MS (ESI, *m/z*): $[M + H]^+$ calculated for $C_{12}H_{23}ClINO_3$, 392.04; found: 392.24.

**_N_-(2-(2-(2-((6-chlorohexyl)oxy)ethoxy)ethoxy)ethyl)-2-iodoacetamide (Probe 2)**

Yellow oil, Yield: 9%.

$^{1}$H NMR (500 MHz, CDCl3) δ 6.87 (d, J = 123.6 Hz, 1H), 3.98 (s, 1H), 3.64 (s, 1H), 3.62 – 3.56 (m, 6H), 3.56 – 3.49 (m, 4H), 3.49 – 3.36 (m, 6H), 1.75 – 1.67 (m, 2H), 1.58 – 1.48 (m, 2H), 1.42 – 1.26 (m, 4H). $^{13}$C NMR (126 MHz, CDCl3) δ 167.85, 71.85, 71.13, 70.97, 70.70, 69.96, 69.34, 45.62, 43.22, 33.11, 30.01, 27.26, 25.99.

MS (ESI, *m/z*): $[M + H]^+$ calculated for $C_{14}H_{27}ClINO4$, 436.07; found: 436.20.

**_N_-(18-chloro-3,6,9,12-tetraoxaoctadecyl)-2-iodoacetamide (Probe 3)**

Yellow oil, Yield: 10%.

$^{1}$H NMR (500 MHz, CDCl3) δ 6.93 (d, J = 71.3 Hz, 1H), 3.64 (s, 1H), 3.63 – 3.49 (m, 14H), 3.49 – 3.42 (m, 3H), 3.42 – 3.36 (m, 3H), 1.75 – 1.67 (m, 2H), 1.57 – 1.48 (m, 2H), 1.42 – 1.34 (m, 2H), 1.34 – 1.26 (m, 2H). $^{13}$C NMR (126 MHz, CDCl3) δ 167.84, 71.71, 71.07, 71.02, 70.99, 70.82, 70.76, 70.54, 69.87, 45.50, 43.11, 33.00, 29.88, 27.15, 25.87.

MS (ESI, *m/z*): $[M + H]^+$ calculated for $C_{16}H_{31}ClINO_5$, 480.09; found: 480.29.

#### Analytical and Biochemical Methods

##### Protein Sequences:

General color code: Hisx6-tag – TEV cleavage site – Protein sequence – Mutation – Catalytic residue

>TrxA C33S

MKHHHHHHPMSDKIIHLTDDSFDTDVLKADGAILVDFWAEWSGPCKMIAPILDEIADEYQGK
LTVAKLNIDQNPGTAPKYGIRGIPTLLLFKNGEVAATKVGALSKGQLKEFLDANLA

>HT7

MHHHHHHHHHHENLYFQ|GIGTGFPFDPHYVEVLGERMHYVDVGPRDGTPVLFLHGNPTSS
YVWRNIIPHVAPTHRCIAPDLIGMGKSDKPDLGYFFDDHVRFMDAFIEALGLEEVVLVIHDW
GSALGFHWAKRNPERVKGIAFMEFIRPIPTWDEWPEFARETFQAFRTTDVGRKLIIDQNVFIE
GTLPMGVVRPLTEVEMDHYREPFLNPVDREPLWRFPNELPIAGEPANIVALVEEYMDWLHQS
PVPKLLFWGTPGVLIPPAEAARLAKSLPNCKAVDIGPGLNLLQEDNPDLIGSEIARWLSTLEI

##### Protein Expression and Purification

###### Thioredoxin A (TrxA$^{C33S}$):[3]

TrxA C33S was cloned, expressed and purified as previously reported (doi: 10.1002/ejoc.202401246 and doi: 10.1021/jacsau.5c00692). In brief, for protein expression the plasmid encoding for His6-TrxA C33S was transformed in *E. coli* BL21 (DE3). Cells obtained from an overnight Lysogeny broth (LB) culture (supplemented with 50 µg/ml Kanamycin) were diluted 1:100 in LB medium (kanamycin) and incubated at 37 °C, 180 rpm until an optical density at 600 nm ($OD_{600}$) of 0.6 was reached. Transgene expression was induced by the addition of 0.5 mM isopropyl-β-D-1-thiogalactopyranoside (IPTG) and cells were grown overnight at 16 °C, 180 rpm. Cells were subsequently harvested by centrifugation (6000 ×g, 20 min).

All purification steps were performed at 4 °C or on ice. The two-step purification procedure of TrxA$^{C33S}$ comprised a His-affinity chromatography followed by a size-exclusion chromatography column (SEC). The cell pellet obtained from a 500 mL expression was resuspended in 40 mL lysis buffer (phosphate-buffered saline (PBS), 20 mM imidazole, pH 7.5) supplemented with DNase I (AppliChem) and complete mini EDTA-free protease inhibitors (Roche). Cells were lysed by homogenization using an

EmulsiFlexC5 (Avestin Inc.) and clear lysate was obtained by centrifugation (24,446 ×g, 30 min, 4 °C) and followed by filtration using a Whatman TM folded filter (Cytiva). The cleared lysate was loaded onto Nickel-NTA agarose beads (Qiagen). After sample loading, the beads were washed with wash buffer (PBS, 20 mM Imidazole, pH 7.5), followed by elution of His6-TrxA (PBS, 500 mM imidazole, pH 8). Elution containing the protein were concentrated using an Amicon Ultracell Centrifugal filter unit (MWCO 3 kDa, Merck Millipore) and applied onto a HiLoad Superdex 200 (10/300) column (Cytiva), equilibrated with PBS, pH 7.5 on an Äkta Pure FPLC system (Cytiva). Protein concentration was measured by Spectrophotometer (DeNovix DS-11, Biozym). Fractions containing the pure protein were pooled and aliquots were stored at -80 °C after being flash-frozen in liquid nitrogen.

The correct size and purity of proteins were assessed by 15% sodium dodecyl sulfate−polyacrylamide gel electrophoresis (SDS−PAGE) and liquid chromatography-mass spectrometry (LC-MS/MS) analysis.

**HaloTag (HT7):[4]**

pET51b-His-TEV-HaloTag7 was a gift from Kai Johnsson (Addgene plasmid #167266) which was chemically transformed in *E. coli* strain BL21(DE3)-pLysS for protein expression. Lysogeny broth (LB) cultures were grown at 37 °C until an $OD_{600}$ of 0.8 was reached. Transgene expression was induced by the addition of 0.5 mM IPTG, and cells were grown overnight at 17 °C in the presence of 1 mM $MgCl_2$. Cells were subsequently harvested by centrifugation (6000 ×g, 20 min).

The cell pellet obtained from a 500 mL expression was resuspended in 40 mL lysis buffer PBS, 20 mM imidazole, pH 7.5) supplemented with DNase I (AppliChem) and complete mini EDTA-free protease inhibitors (Roche). Cells were lysed by homogenization using an EmulsiFlexC5 (Avestin Inc.) and clear lysate was obtained by centrifugation (24,446 ×g, 30 min, 4 °C) and followed by filtration using a Whatman TM folded filter (Cytiva). The cleared lysate was loaded onto Nickel-NTA agarose beads (Qiagen). After sample loading, the beads were washed with wash buffer (PBS, 20 mM Imidazole, pH 7.5), followed by elution of His6-TEV-HaloTag (PBS, 500 mM imidazole, pH 8). Elution containing the protein were concentrated using an Amicon Ultracell Centrifugal filter unit (MWCO 3 kDa, Merck Millipore). Buffer was exchanged using a PD-10 desalting column (Cytiva) for PBS, 150 mM NaCl (pH 7.5). HT7 concentration was measured by Spectrophotometer (DeNovix DS-11, Biozym). Fractions containing the pure protein were pooled and aliquots were stored at -80 °C after being flash-frozen in liquid nitrogen.

The correct size and purity of proteins were assessed by 7.5% SDS−PAGE.

##### *in-vitro* Probe Treatment

100 µM TrxA$^{C33S}$ in PBS buffer (100 µL) was treated with respective probe **1**, **2** or **3** (1 mM, TrxA:probe 1:10 ratio) or DMSO as control for 2 hours at 25° C, 600 rpm. After incubation, the protein was precipitated by adding 4x excess amount of cold acetone at -20° C overnight. Proteins were pelleted by centrifugation at 4° C, 13,000 x g for 10 min. The supernatants were then removed, and the protein pellets were washed twice with cold methanol, reconstituted with PBS buffer and sonicated with a rod sonicator (10 s, 20% intensity). The modified and unmodified TrxA$^{C33S}$ were assessed by 15 % SDS-PAGE gel and LC-MS/MS intact protein measurement.

##### HaloTag Labeling

In a solution of activity buffer (50 mM HEPES, 150 mM NaCl, pH 7.3, 50 µL), TrxA$^{C33S}$ was labelled with HaloTag protein for 15 min at 25° C, 450 rpm. Labelled proteins were then prepared for SDS-PAGE analysis.

##### Pulse-chase Assay

After the HaloTag labeling, proteins were treated with HaloTag-TMR fluorescent ligand (1 µM, MedChemExpress, #HY-D2270) for another 15 min at 25° C, 450rpm. Treated proteins were then prepared for SDS-PAGE analysis, as described below.

##### Cell Culture

HEK293T, Hela or SH-SY5Y cells were cultured in Dulbeccos Modified Eagles Medium – high glucose (DMEM, Merck) supplemented with 10% fetal bovine serum (FBS, Merck) and 2 mM stable L-glutamine (Merck) at 37 °C and 5% $CO_2$ atmosphere.

Cells were harvested when they were grown to 90 – 95% confluence. The cell culture medium was firstly removed, then the cells were washed twice with cold PBS. Cells were then scraped in 1 mL PBS and transferred to a 1.5 mL Eppendorf tube. Afterwards the cell suspension was centrifuged at 1000 rpm, 4 ºC for 4 minutes. The supernatant was then carefully removed, and the cell pellet was stored at -80°C until further use.

##### Cell Lysis

Cells were lysed with 200 – 400 µL of lysis buffer (1% NP40, 0.2% SDS in 25 mM Hepes, 7.5 pH) via rod sonication (10 s, 20% intensity). The lysates were clarified by centrifugation (13,000 xg, 4 °C, 10 min) and the supernatant was transferred into a new Eppendorf tube. The lysates were stored at -80 °C or processed directly.

##### Protein concentration measurement

Protein concentration measurement was performed with a Pierce™ BCA Protein Assay Kit (Thermo Scientific).

##### HaloTag Labeling in Cell Lysates

100 µg protein lysates were diluted in activity buffer, to total volume of 50 µL before supplementing with modified or unmodified $TrxA^{C33S}$. HaloTag protein was incorporated and incubated for 15 min at 25° C, 450rpm.

##### In-gel Shift Assay

Before loading onto the respective 7.5%, 10% or 15% polyacrylamide gels, 20 µL protein solution was mixed with 5 µL of 5× Laemmli buffer (10% [w/v] SDS, 50% [v/v] glycerol, 25% [v/v] β-mercaptoethanol, 0.5% [w/v] bromophenol blue, 315 mM Tris–HCl, 100 mM DTT, pH 6.8), boiled for 5 min at 95° C, loaded and ran with 150 V for 1 h in 1x running buffer (25 mM Tris, 0.192 M glycine, 0.1% SDS). As a reference, two types of protein markers were used: Color Pre-stained Protein Standard, Broad Range (10-250 kDa) (New England Biolabs GmbH), BenchMark™ Fluorescent Protein Standard (Invitrogen™). For loading control, proteins were stained with a Coomassie staining solution (0.25% Coomassie Blue R-250, 10% acetic acid, 50% MeOH) and properly destained with destaining solution (20% MeOH, 10% acetic acid). Afterwards, the gel was scanned on Amersham ImageQuant 800 (Cytiva).

##### Western Blot

After separating the proteins on SDS-PAGE, they were blotted on a PVDF membrane using a Trans-Blot Semi-Dry Transfer System (Bio-Rad). For this, the SDS-Gel and filter paper were equilibrated in transfer buffer (48 mM Tris, 39 mM glycine, 0.0375% [m/v] SDS, and 20% [v/v] methanol) for 5 min at room temperature. The PVDF membrane was first activated for 1 min in MeOH and then equilibrated in transfer buffer for 5 min at room temperature. After preparing the blotting sandwich, the protein transfer was carried out for 30 min at 25 V. Afterwards, the membrane was blocked in 10 mL of blocking buffer (5% milk solution in PBST (PBS + 0.5% Tween)) for 1 h. Subsequently, 10 mL of anti-

TrxA antibody (1:1000, Antibodies Online, #ABIN7141748) diluted in blocking solution was added, and the mixture was incubated for overnight at 4 °C. The membrane was washed three times for 10 min with PBST before 1 µL of the anti-rabbit (1:10000) secondary antibody diluted in 10 mL of blocking solution was added. After 1 h of incubation at room temperature, the membrane was washed three times for 10 min with PBST while shaking at room temperature. Similarly, after stripping the membrane, it was incubated with anti-HaloTag antibody (1:1000, Promega, # G928A) for overnight at 4 °C and again with anti-rabbit secondary antibody for 1 h with PBST washes. For imaging, a 1:1 mixture composed of ECL detection reagents (Cytiva) was prepared and added to the membrane for chemiluminescence signal detection. Finally, images of the WB were taken by developing using the Amersham ImageQuant 800 (Cytiva).

**Direct Injection for Intact Protein Measurement**

For top-down measurements, purified proteins (10 µM) were desalted on a ZipTip with C4 resin (Merck-Millipore, ZTC04S096). The resin with loaded protein was washed thrice with a buffer containing 1 % (v/v) formic acid and the protein was eluted with a buffer (300 µL) containing 50 % (v/v) acetonitrile and 0.5 % (v/v) formic acid. For the ESI-MS measurements, an Orbitrap Eclipse Tribrid Mass Spectrometer (Thermo Fisher Scientific, Waltham, USA) was used via direct injection with a HESI-Spray source and FAIMS interface in a positive, intact protein mode. The FAIMS compensation voltage (CV) was searched by a continuous scan. The most intense signal was obtained at -27 CV for TrxA C33S and 24 CV for HaloTag. The MS spectra were acquired with 120,000 FWHM, AGC target 100 and 2 microscans. Spectra were deconvoluted and processed further with UniDec 8.1.0[5].

**Immunoprecipitation of HaloTag Protein**

Immunoprecipitation was done with ChromoTek Halo-Trap Magnetic Agarose beads (ChromoTek, #otma) according to manufacturers' recommendation with the following changes. 100 µg of HEK293T cell lysates were treated with 200 nM of HaloTag and 2 µM of modified TrxA C33S for 15 min at 25° C, 450 rpm and 2 µL of beads were used for binding step, eluted beads with 20 µL of 2x Laemmli buffer and boiled the samples at 95° C for 5 min before loading onto a gel. Input and bound fractions were analyzed by SDS-PAGE and Western blot as described above.

**On-bead digest for mass-spectrometry**

Following the manufacturer's protocol, after protein binding, the beads were washed 3x with ice-cold lysis buffer (10 mM Tris/Cl, pH 7.5; 150 mM NaCl; 0.5 mM EDTA; 0.5% NP-40) by centrifugation (2500 x g for 2 min at 4° C) and on magnet. Then the beads were washed 2x with ice-cold wash buffer (10 mM Tris/Cl, pH 7.5; 150 mM NaCl; 0.5 mM EDTA) by centrifugation (2500 x g for 2 min at 4° C) and on magnet. Afterwards, the beads were digested with freshly prepared elution buffer I (50 mM Tris/Cl, pH 7.5; 2 M Urea; 5 µg/µL sequencing-grade Trypsin; 1 mM DTT) for 30 min, 400 rpm at 30° C. After centrifugation (2500 x g for 2 min at 4° C), the peptides were transferred into a new Eppendorf tube. Again, the peptides were eluted from beads with elution buffer II (50 mM Tris/Cl, pH 7.5; 2 M Urea; 5 mM iodoacetamide), centrifuged and combined with previous fraction. The peptides were digested overnight at 32° C, 400 rpm overnight. Next day, the reaction was stopped by the addition of 1 µL formic acid (FA).

To desalt the samples, Sep-Pak C18 cartridges were used. The cartridge was flushed with 1 mL of ACN and 1 mL of ACN and FA mixture (80% ACN + 0.5% FA in MS-$H_2O$). Equilibration was performed three times with 1 mL FA (0.5% in MS-$H_2O$). Afterwards, the sample was loaded into the cartridge. The column was then washed three times with 1 mL of FA (0.5% in MS-$H_2O$). Peptides were eluted twice with 250 µL of ACN and FA mixture. The samples were dried in a SpeedVac. Dry peptides were then reconstituted in 30 µL FA (1% in MS-$H_2O$), vortexed, and placed in a sonication bath for 15 min. Samples were then spun down and transferred into MS vials.

**MS Sample Preparation for Whole Proteome Analysis**

HEK293T cell lysates were prepared under standard conditions using the standard treatment and harvesting procedures in biological triplicates. The volume of a lysate containing 10 µg of proteins was normalized to 10 µL with lysis buffer. A mixture of hydrophilic and hydrophobic carboxylate-coated magnetic beads (10 µg/µL, Cytiva) were taken into a 96-well plate and washed three times with 500 µL of MS-grade $H_2O$. After treating the lysates with modified TrxA$^{C33S}$ and HaloTag, they were added to the beads and thoroughly mixed. 20 µL of 0.25% FA in ACN solution is added in each sample and incubated for 8 min at room temperature. Following the automated SP3 protocol in MicroLab Prep pipetting robot (Hamilton), the samples were incubated further for 2 min with agitation. The plate was then placed onto a magnet to remove supernatant-containing unbound components. After removing the supernatant, the beads were washed two times with 180 µL of 80% EtOH in MS-$H_2O$ and once with 180 µL of ACN and incubated for 30 s at room temperature with agitation. The beads were air-dried after aspirating supernatant. Manually, on-beads digestion was performed to cleave the proteins. In brief, beads were reconstituted in 20 µL of 100 mM ammonium bicarbonate (ABC) buffer and 1 µL of sequencing-grade trypsin (0.5 mg/ml; Promega) and incubated overnight at 37°C while shaking at 650 rpm. Next day, the digested peptides were recovered into 1.5 mL Eppendorf tubes, and the beads were washed once with 79 µL of MS-$H_2O$ after incubating for 5 min at 40 °C while shaking at 850 rpm. Then the beads were washed once with 11.4 µL of 1% FA in MS-$H_2O$. The peptides were collected in the initial 1.5 mL tube while being attached onto the magnet. Finally, the supernatant was transferred to an MS vial and subjected to LC–MS/MS analysis.

**LC-MS/MS Measurement**

MS measurements were performed on a Orbitrap Eclipse Tribrid Mass Spectrometer (Thermo Fisher Scientific) coupled to an UltiMate 3000 Nano-HPLC (Thermo Fisher Scientific) via a Nanospray Flex (Thermo Fisher Scientific) and FAIMS interface (Thermo Fisher Scientific). First, peptides were loaded on an Acclaim PepMap 100 µ-precolumn cartridge (5 µm, 100 Å; 300 µM ID x 5 mm, Thermo Fisher Scientific). Then, peptides were separated at 40 °C on a PicoTip emitter (noncoated, 15 cm, 75 µm ID, 8 µm tip, New Objective) that was in house packed with Reprosil-Pur 120 C18-AQ material (1.9 µm, 150 Å, Dr. A. Maisch GmbH). The LC buffers consiststed of MS-grade $H_2O$ (A) and ACN (B) both supplemented with 0.1% formic acid. The short gradient was run from 4-35.2% B during a 60 min method (0-5 min 4%, 5-6 min to 7%, 7-36 min to 24.8%, 37-41 min to 35.2%, 42-46 min 80%, 47-60 min 4%) at a flow rate of 300 nL/min.

**Data-independent acquisition**

FAIMS was performed with one CV at -45 V. One DIA cycle comprised of one $MS^1$ scan followed by 30 $MS^2$ scans. The mass spectrometer was operated in DIA mode with following settings: Polarity: positive; $MS^1$ Orbitrap resolution: 60k; $MS^1$ AGC target: standard; $MS^1$ maximum injection time: 50 ms; $MS^1$ scan range: m/z 200-1800; RF Lens: 30%; Precursor Mass Range: m/z 500-740; isolation window: m/z 4; window overlap: m/z 2; $MS^2$ Orbitrap resolution: 30k; $MS^2$ AGC target: 200%; $MS^2$ maximum injection time: auto; HCD collision energy: 35%; RF Lens: 30%; $MS^2$ scan range: auto.

**Data-Independent Acquisition-Neural Network (DIA-NN)**

Thermo ∗.raw files were analyzed with DIA-NN 2.1.0[6], and peptides were searched against the UniProt database for *Homo sapiens* (taxon identifier: 9606) with included contaminants and decoys (193632 entries in total). The DIA-neural network (NN) settings were as follows: FASTA digest for library-free search/library generation: enabled; Deep learning–based spectra, retention times, and ion mobility prediction: enabled; missed cleavages: 1; maximum number of variable modifications: 1; modifications: N-term M excision; carbamidomethylation, Peptide length range: 7 to 30; Precursor charge range: 2 to

6; precursor range: *m/z* 500 to 740; fragment ion range: *m/z* 200 to 1800; precursor and protein false discovery rate (FDR) level: 1%; match between runs: enabled; library generation: smart profiling; quantification strategy: Robust LC; mass accuracy: 0; and scan windows: 0.

**Processing of quantified data**

For statistical analysis, the "report.pg_matrix.tsv" table was used in Perseus 1.6.14.0.21 (Max Planck Institute of Biochemistry). Then, quantified values were $\log_2$-transformed and columns were assigned with either Control or HaloTag or TrxA-HaloTag treated. Subsequently, the groups were filtered based on the experimental background (at least two valid values out of three columns in at least one group or at least two valid values out of three columns in each group). Further, missing values were replaced from a normal distribution. $-\log_{10}$($p$ values) were obtained by a two-sided one-sample Student's $t$ test over replicates with the initial significance level $\alpha = 0.05$ adjustment by the multiple testing correction method of Benjamini and Hochberg (FDR = 0.05) using the volcano plot function. Finally, the volcano plot values from Perseus software (1.6.10.43) were transferred to GraphPad Prism and visualized accordingly.

#### NMR Spectra

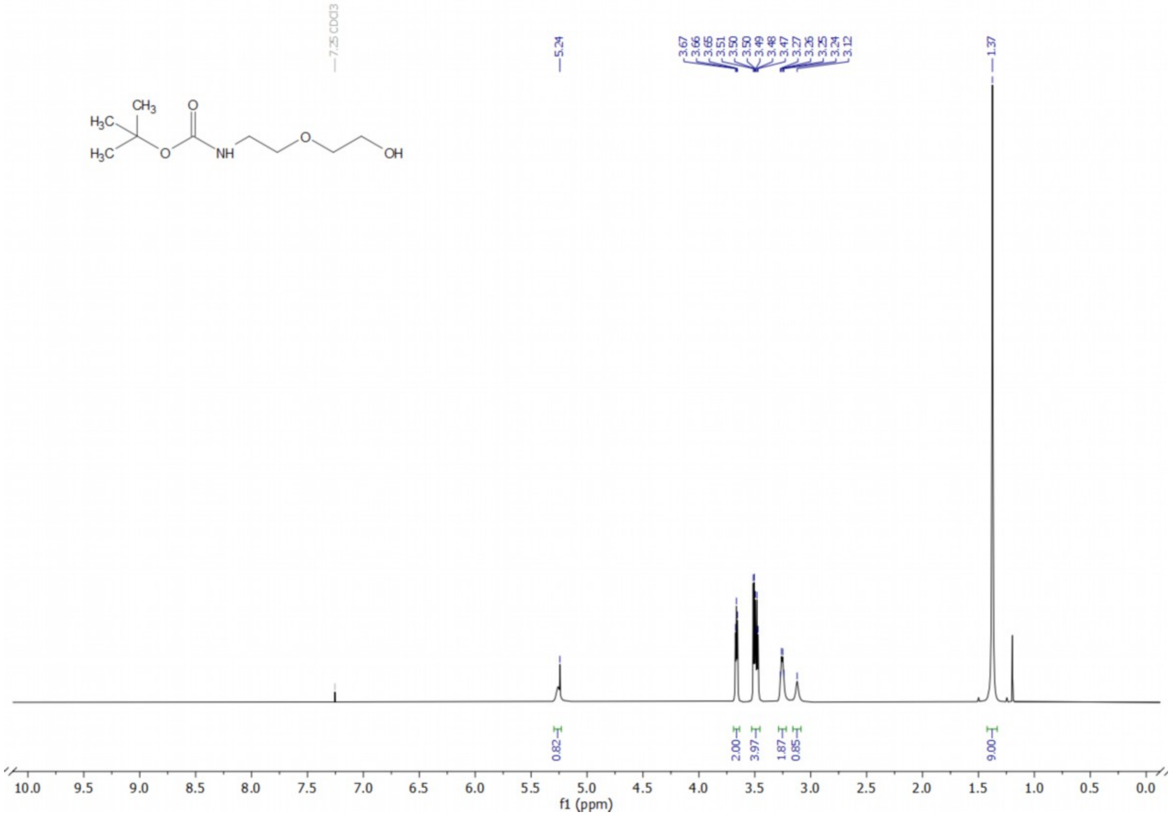

**[1]H NMR of compound 5a**

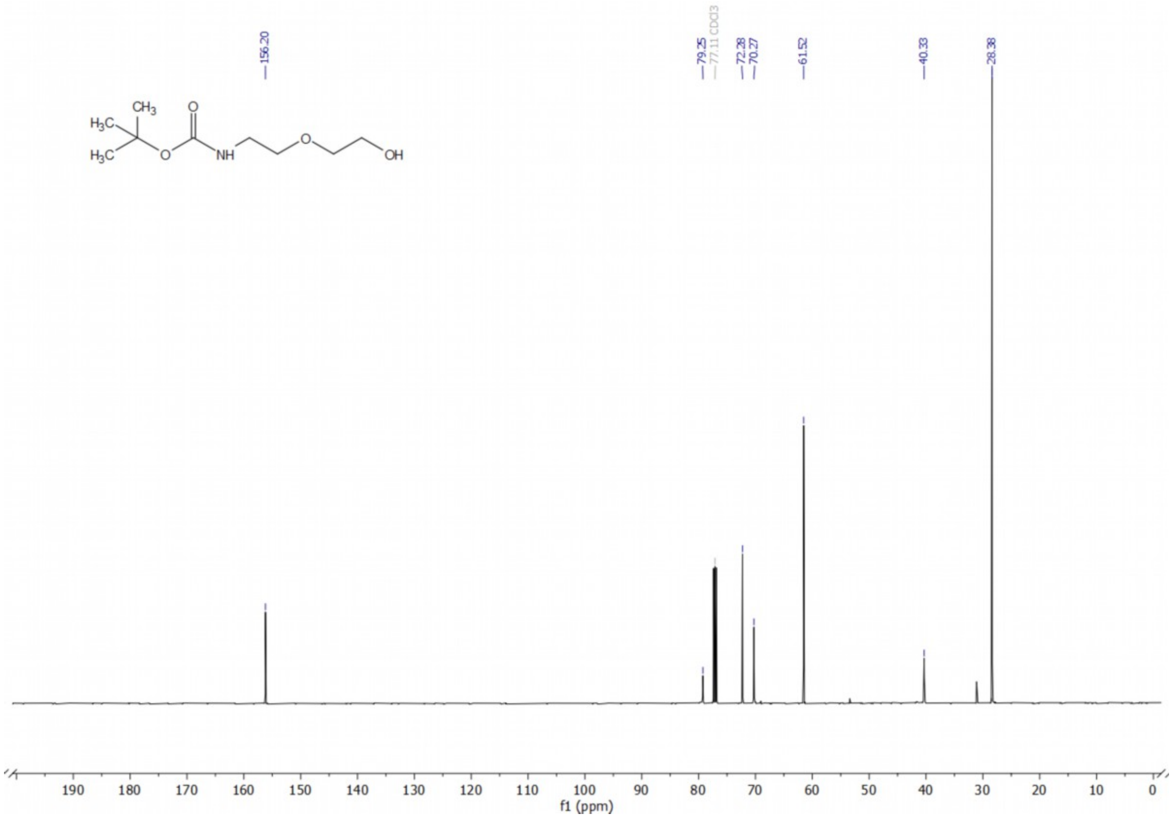

**[13]C NMR of compound 5a**

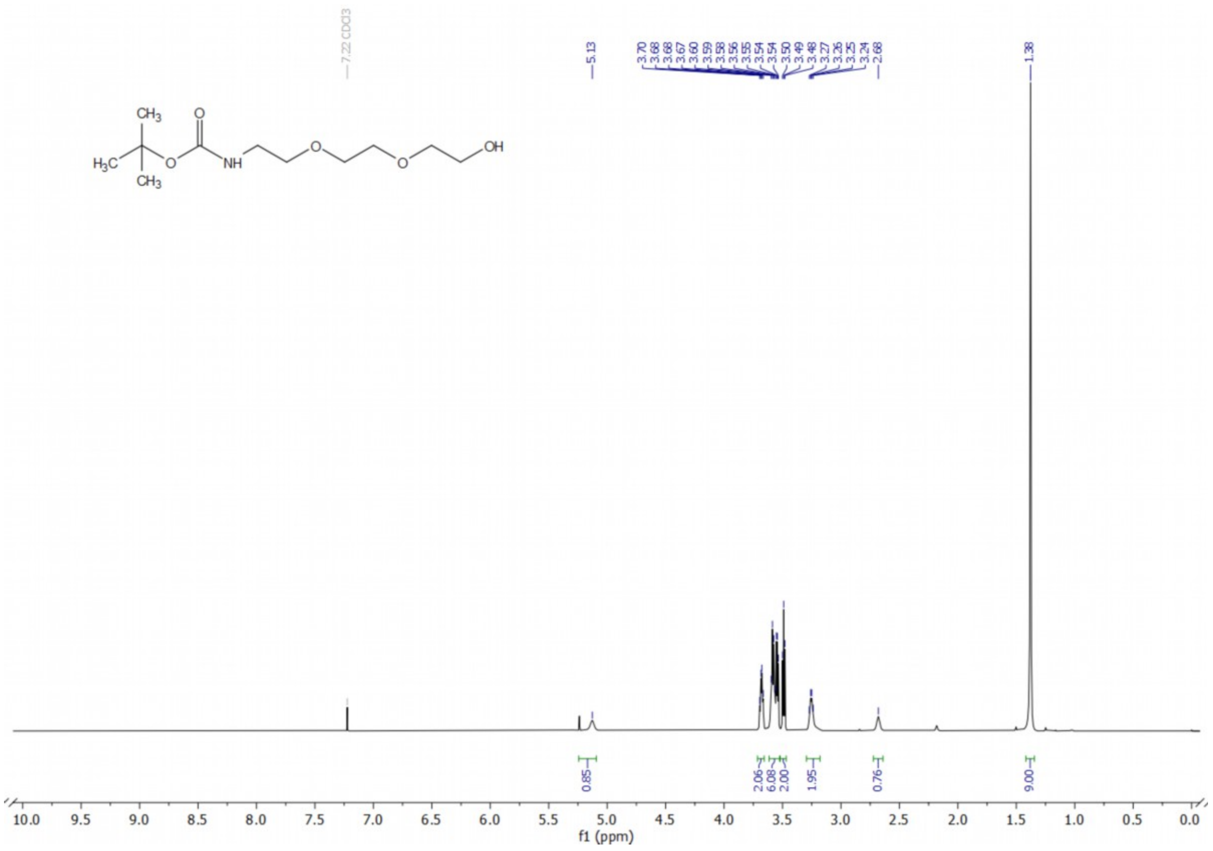

**$^{1}$H NMR of compound 5b**

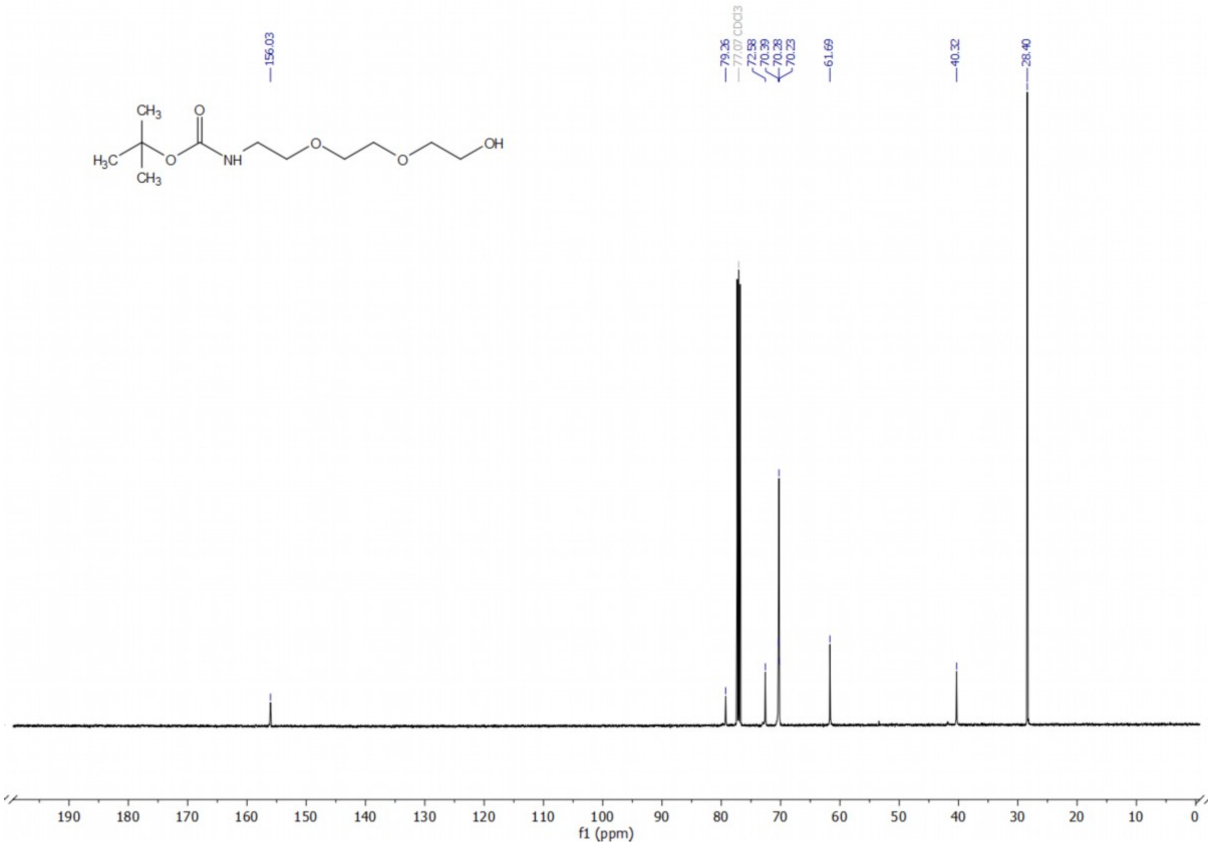

**$^{13}$C NMR of compound 5b**

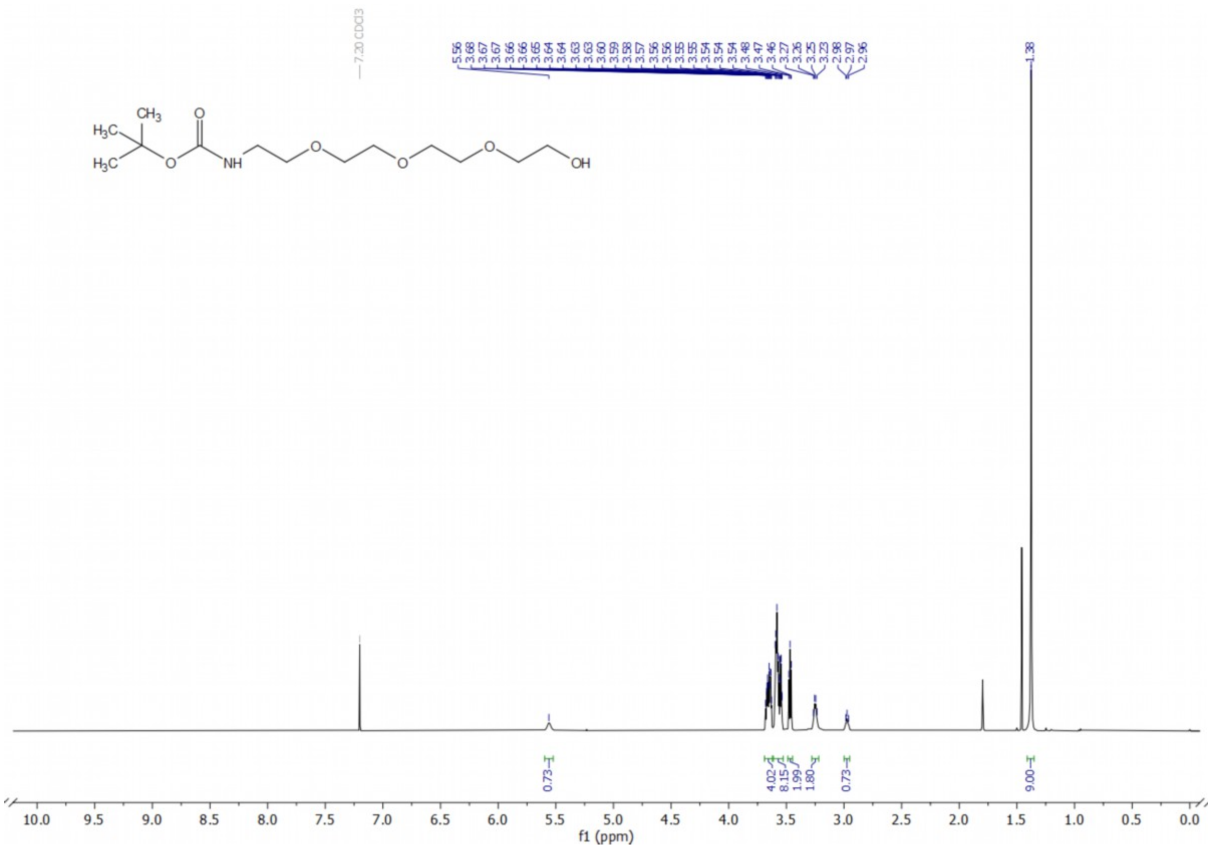

**$^{1}$H NMR of compound 5c**

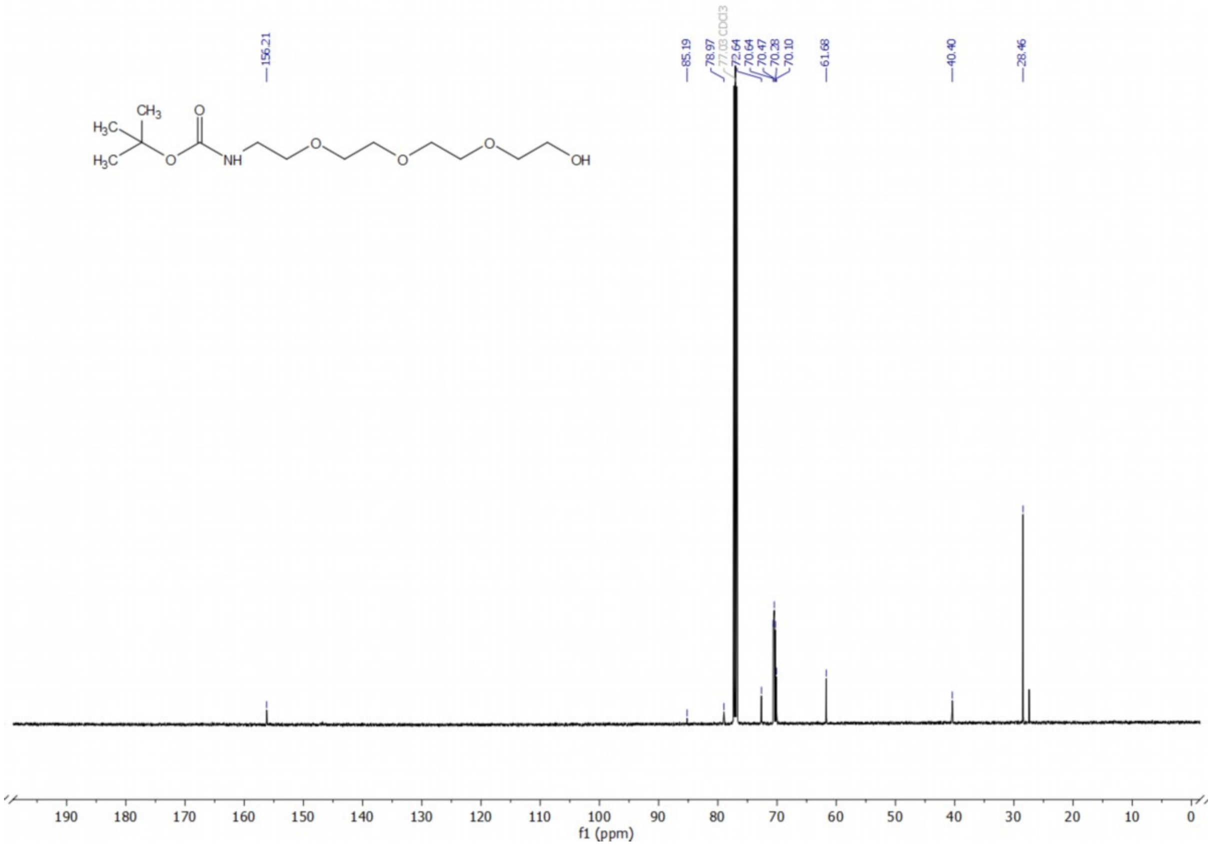

**$^{13}$C NMR of compound 5c**

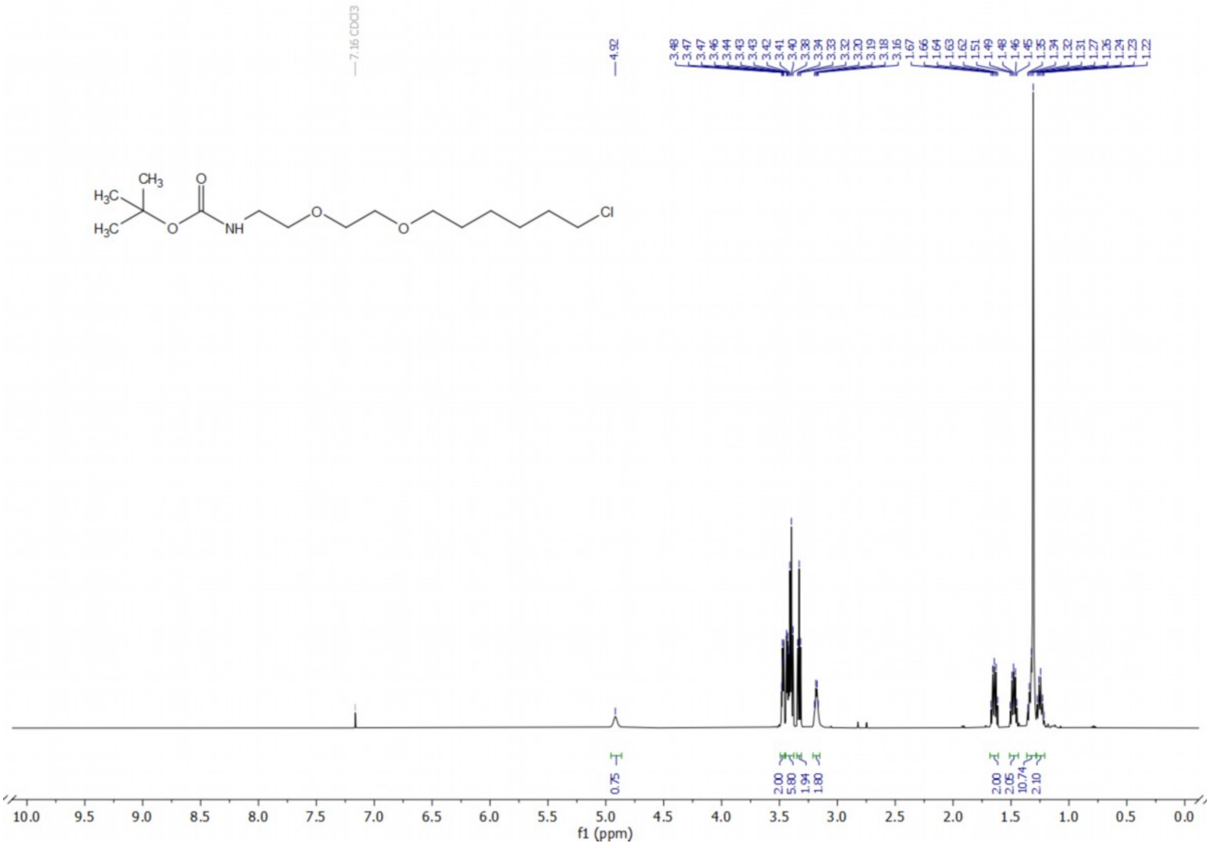

**[1]H NMR of compound 6a**

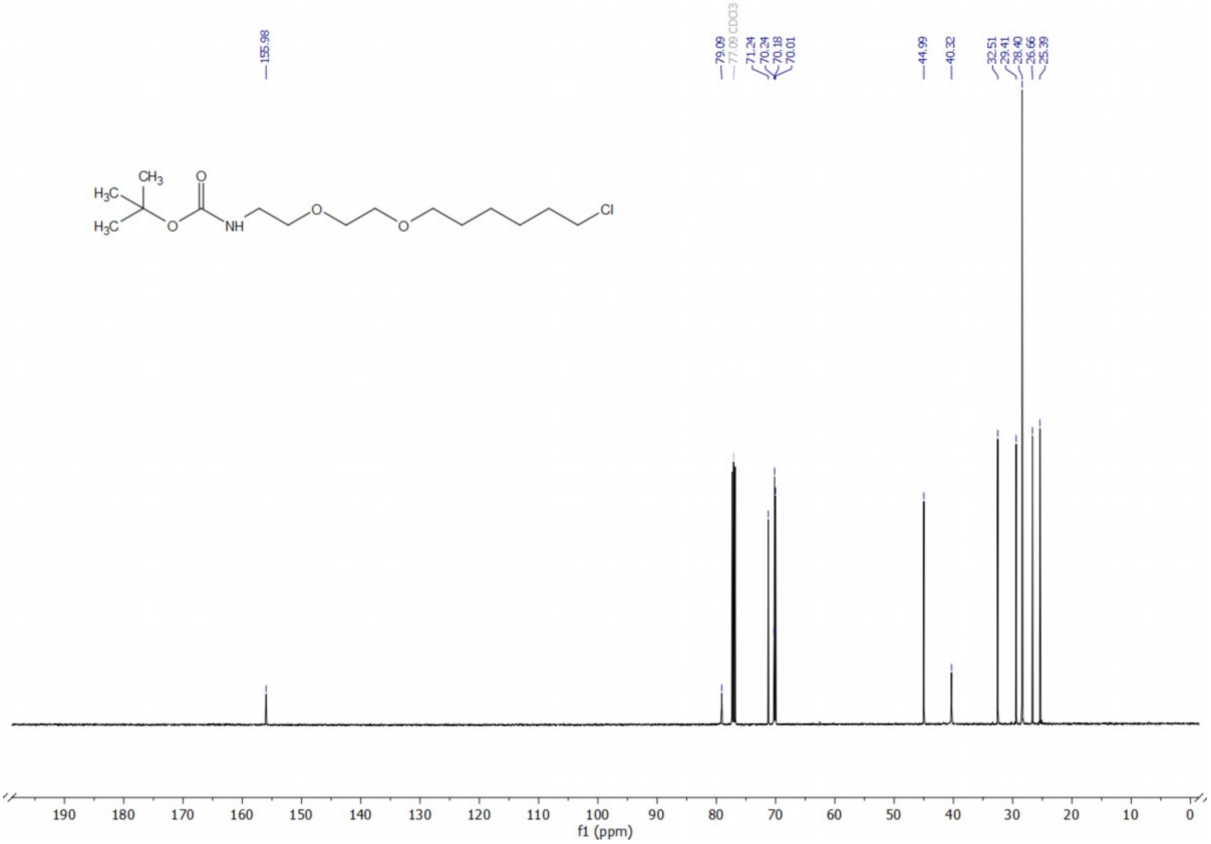

**[13]C NMR of compound 6a**

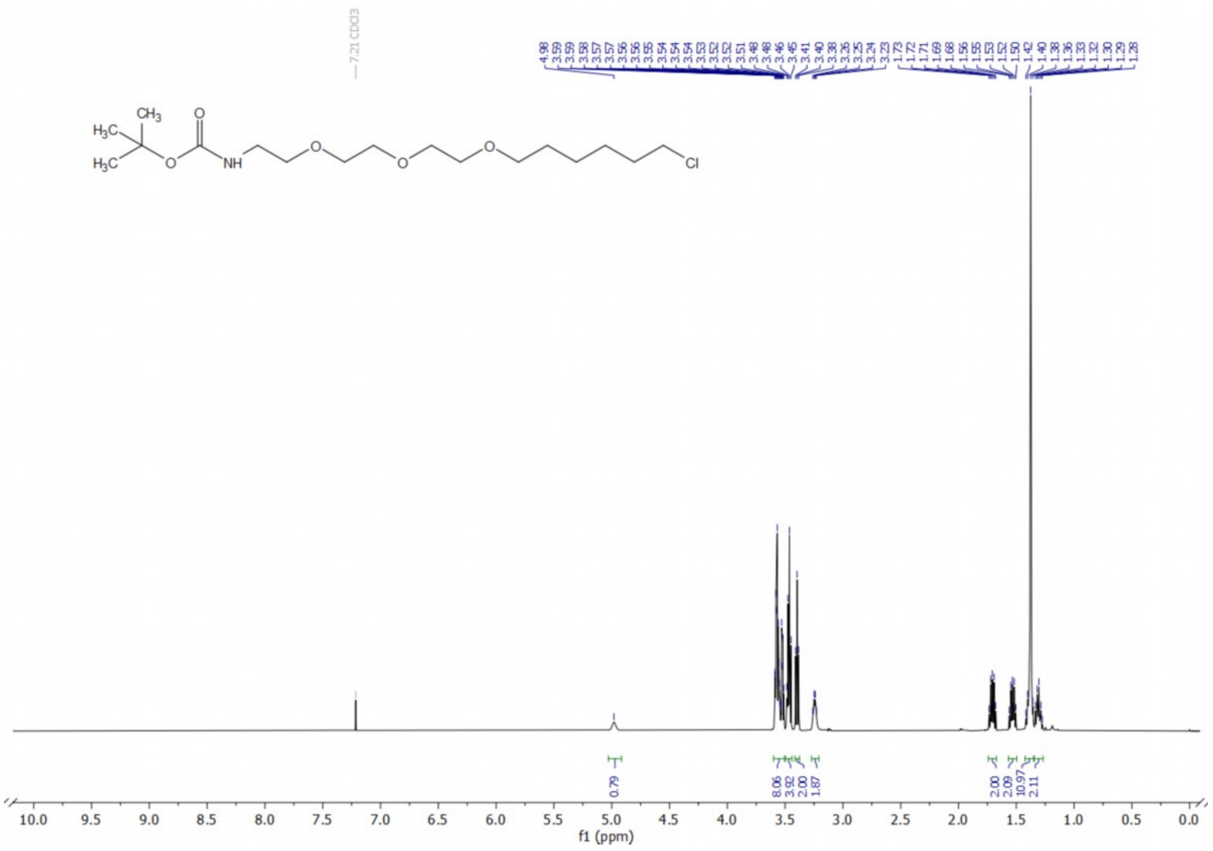

**¹H NMR of compound 6b**

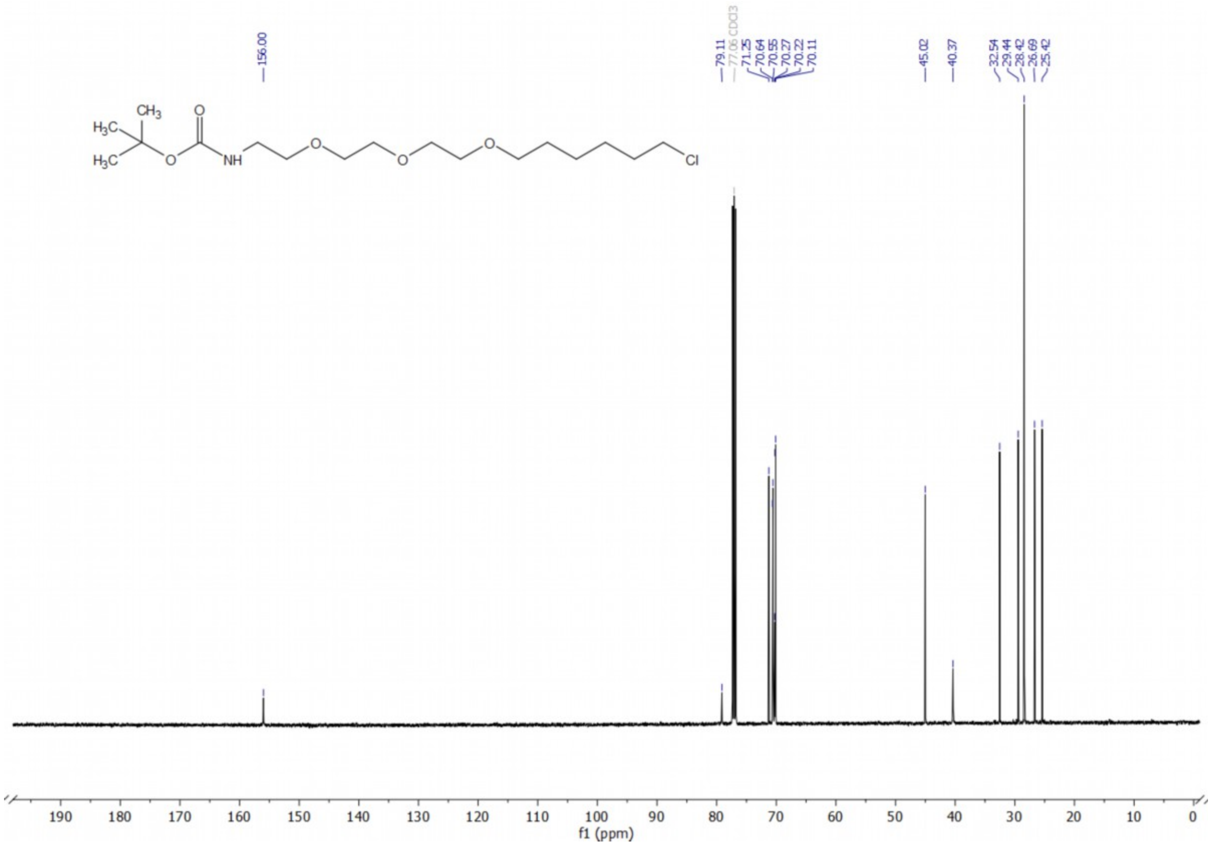

**¹³C NMR of compound 6b**

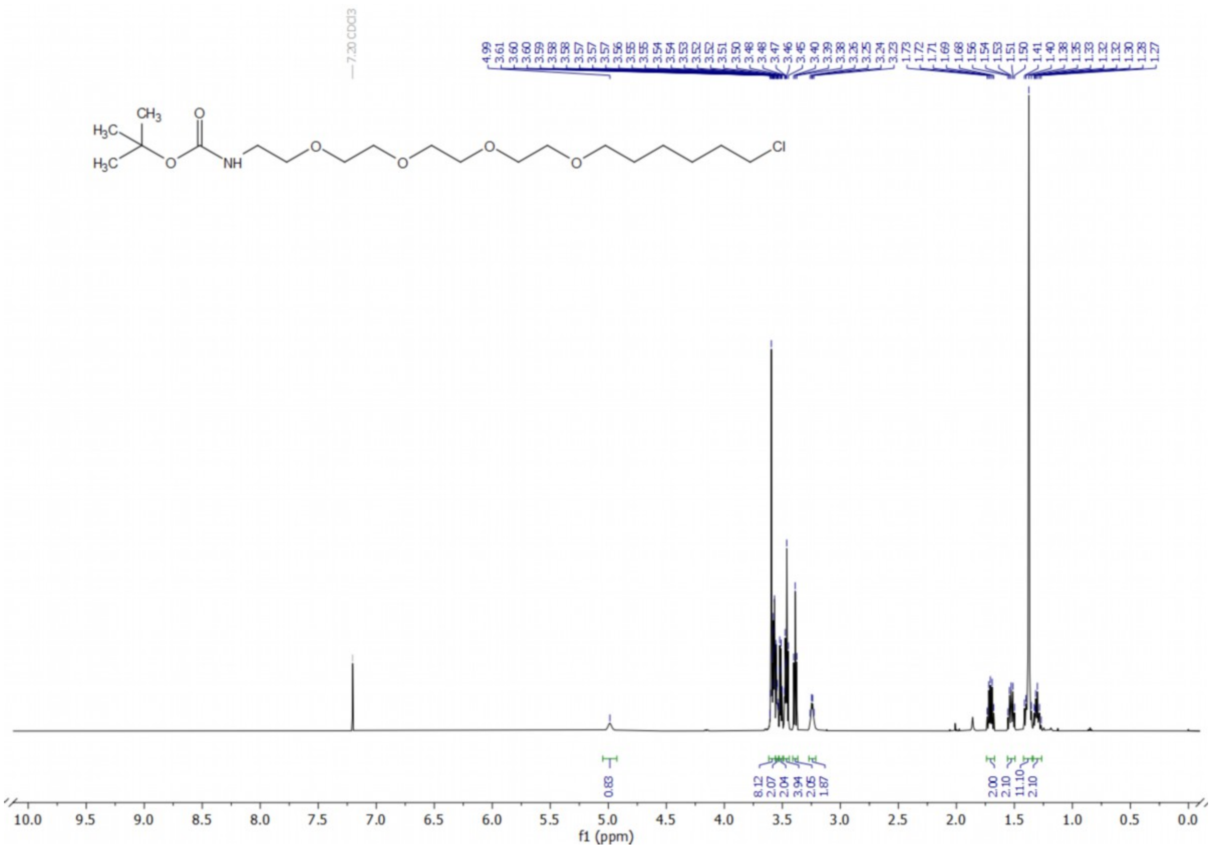

**[1]H NMR of compound 6c**

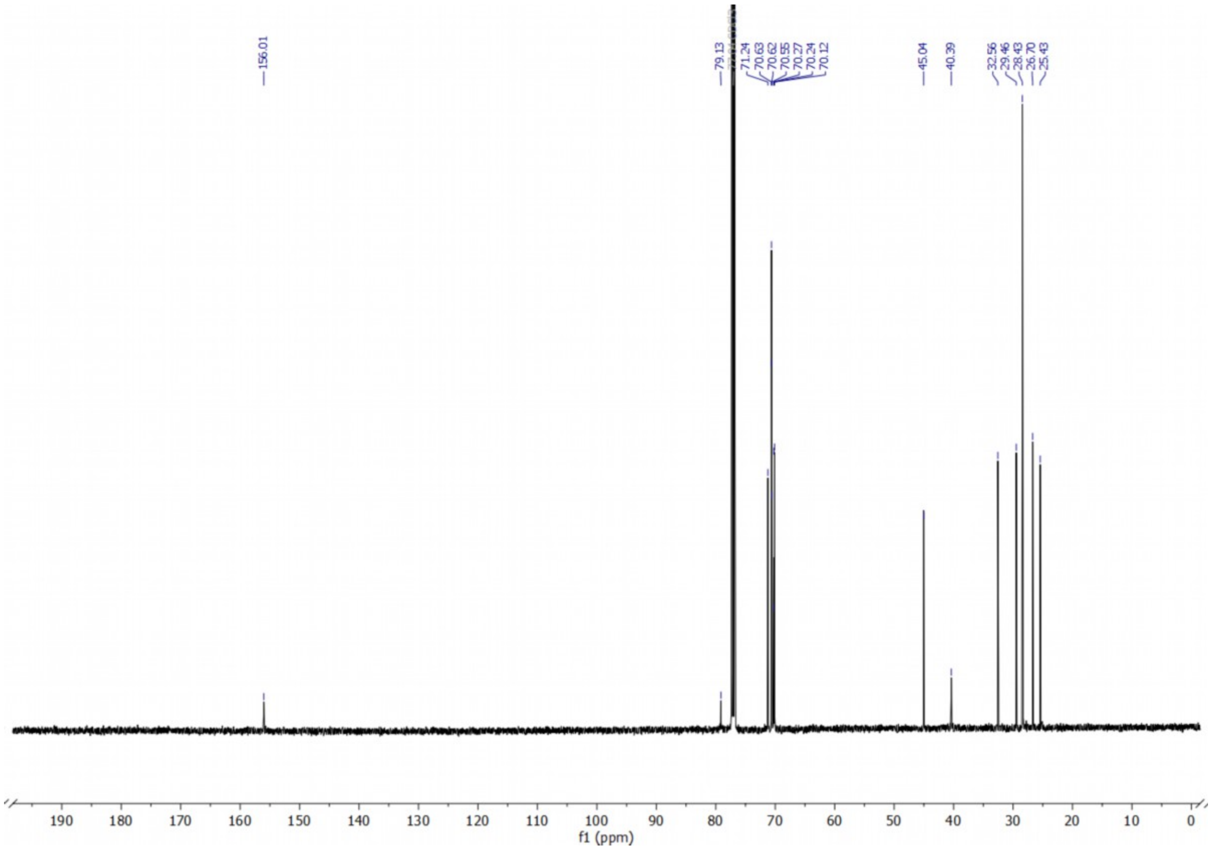

**[13]C NMR of compound 6c**

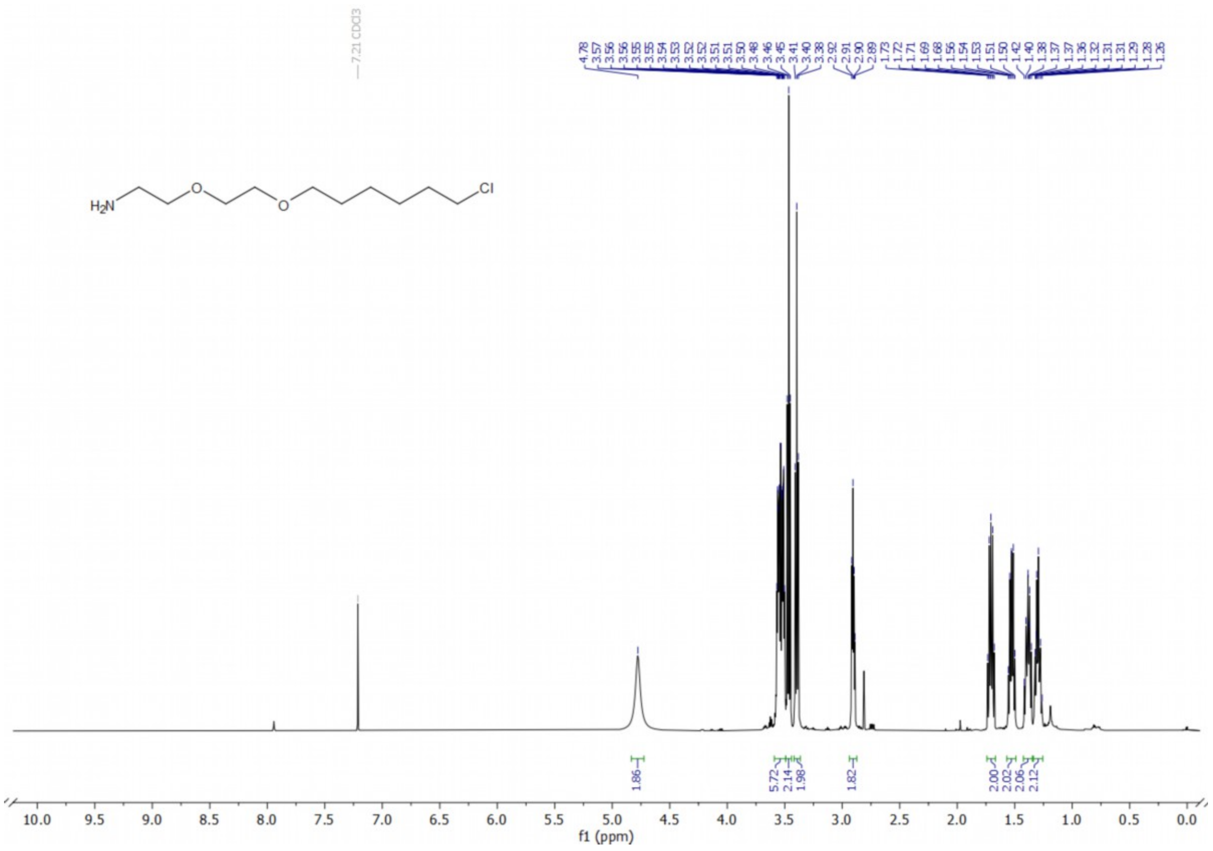

**[1]H NMR of compound 7a**

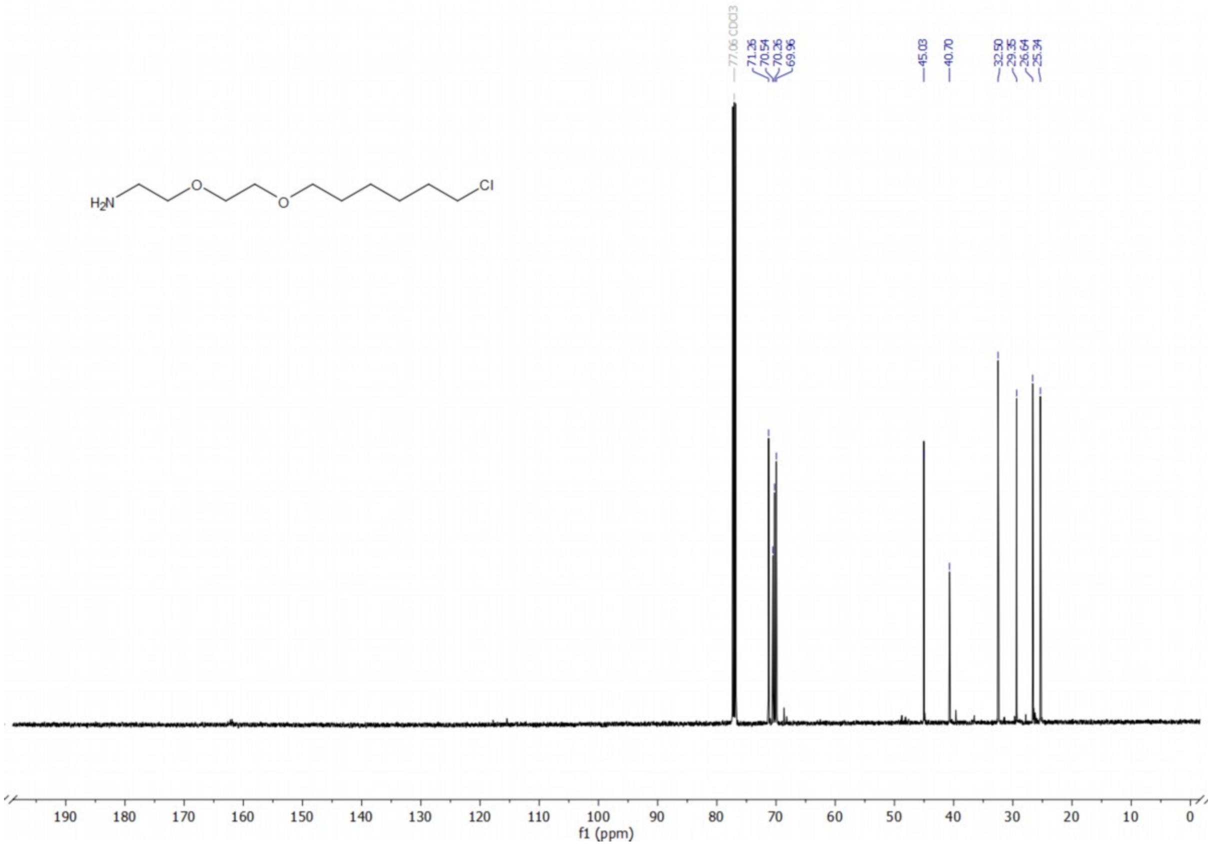

**[13]C NMR of compound 7a**

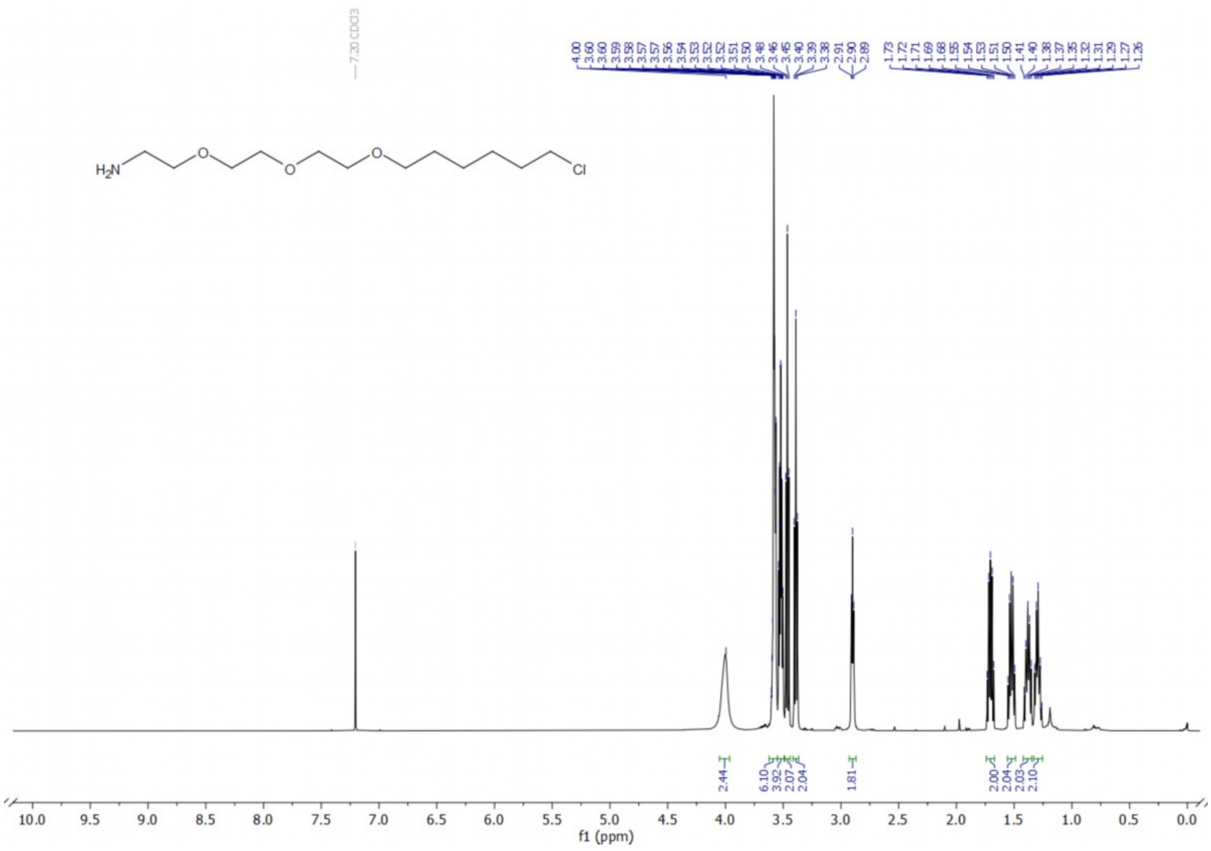

**[1]H NMR of compound 7b**

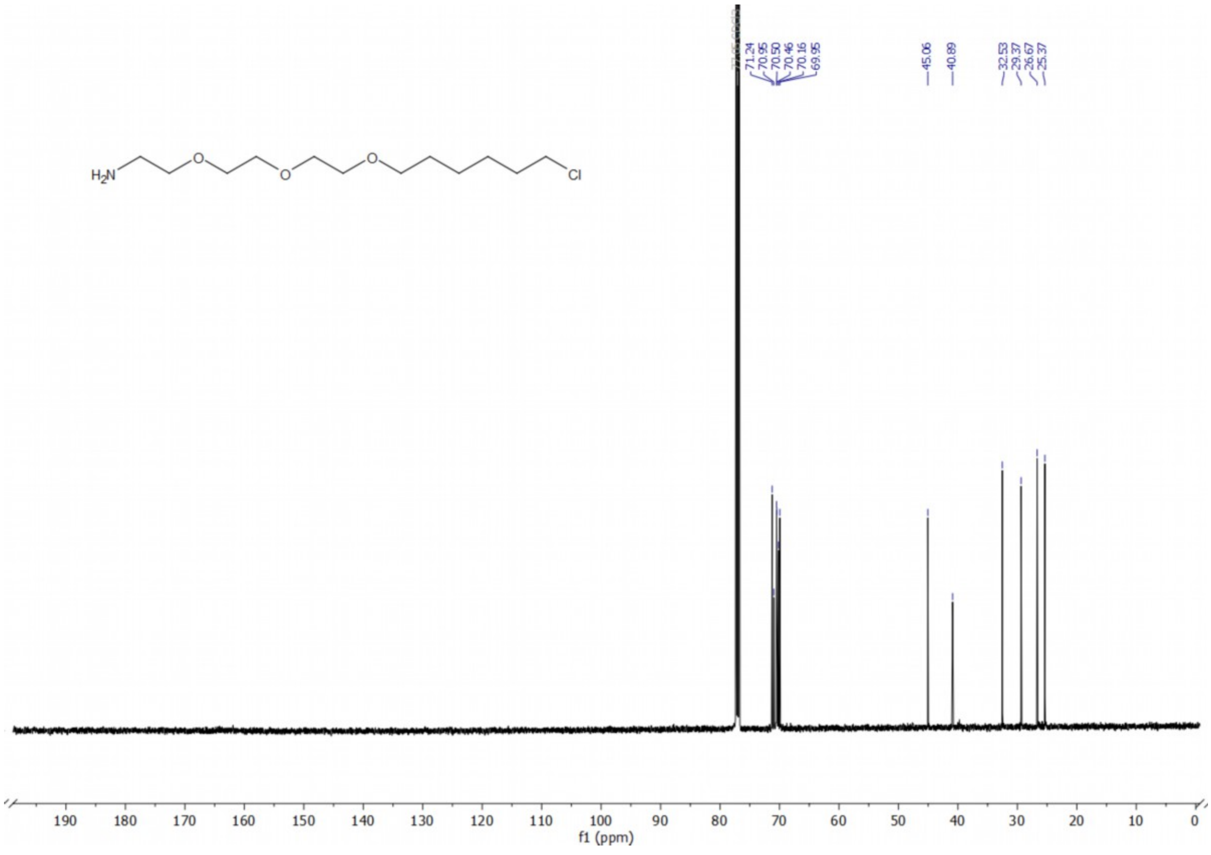

**[13]C NMR of compound 7b**

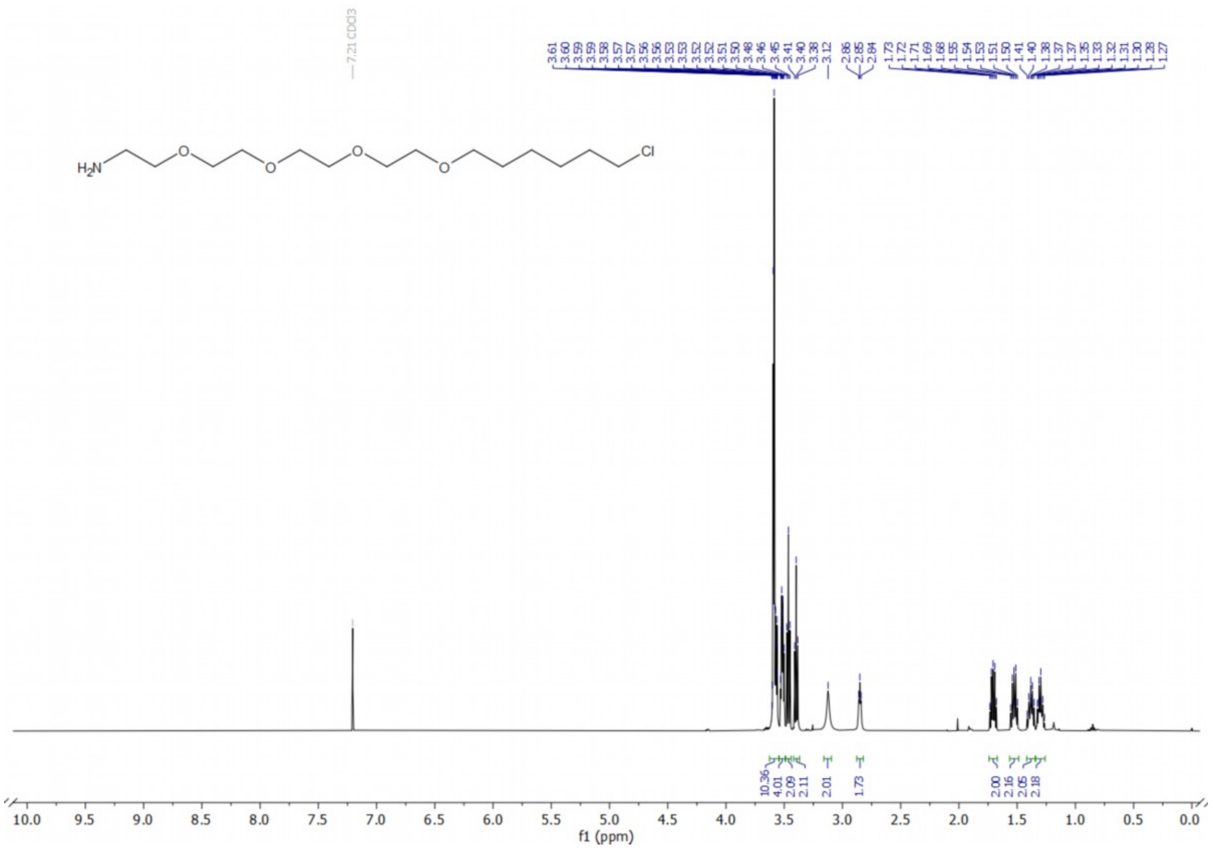

**[1]H NMR of compound 7c**

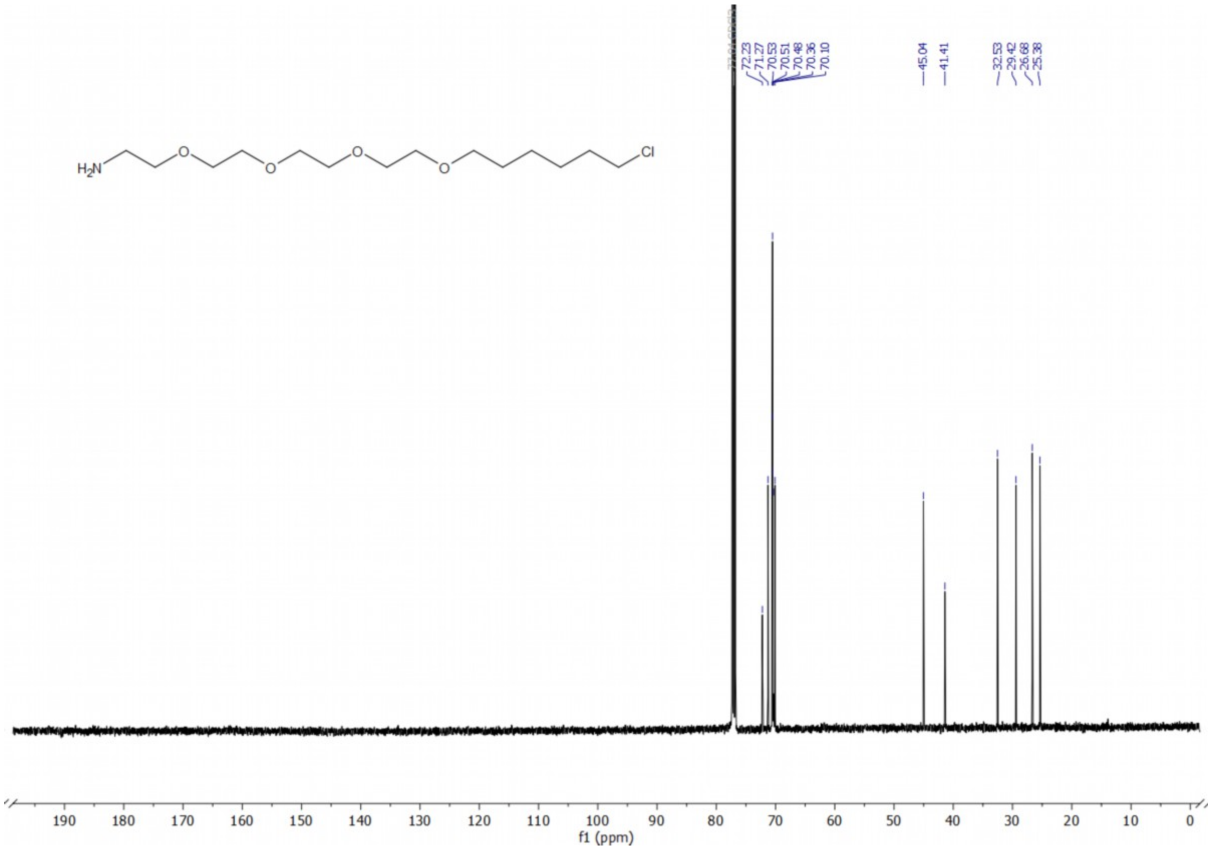

**[13]C NMR of compound 7c**

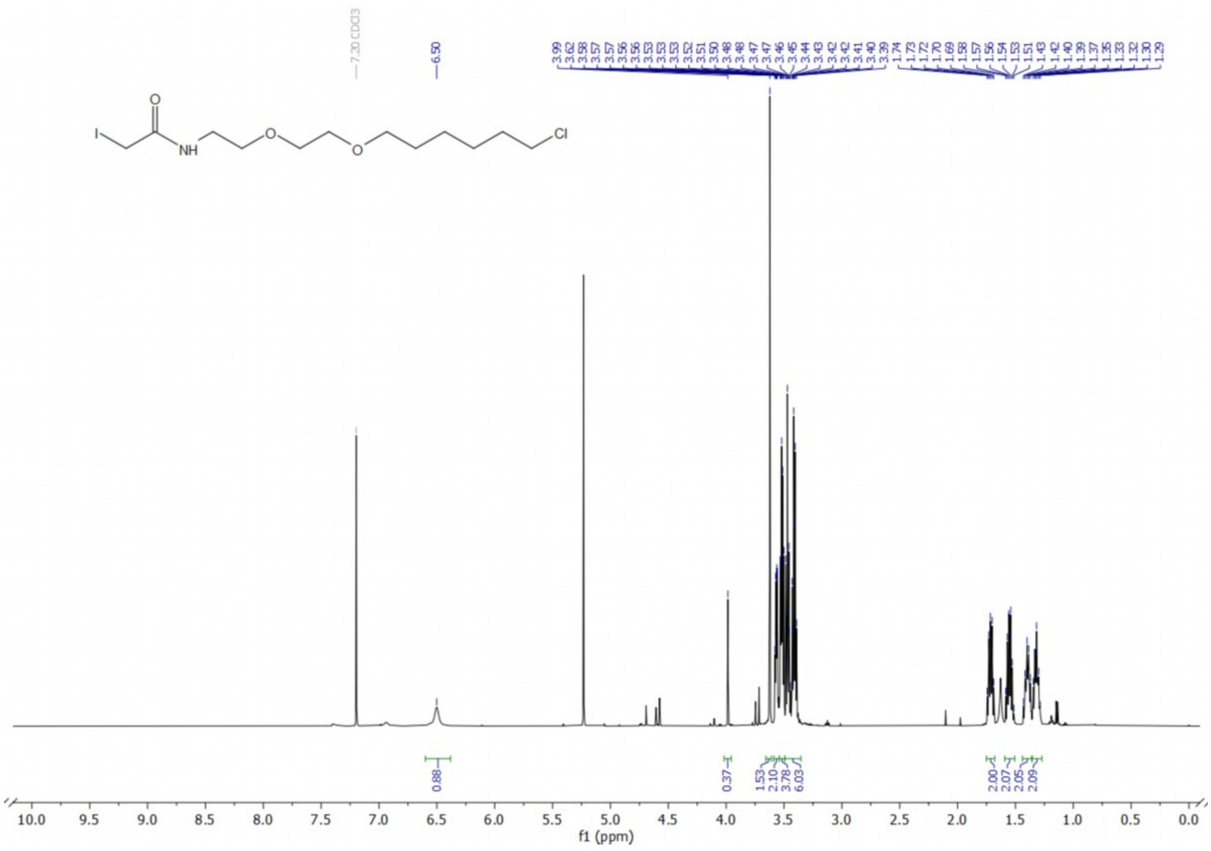

### [1]H NMR of probe 1

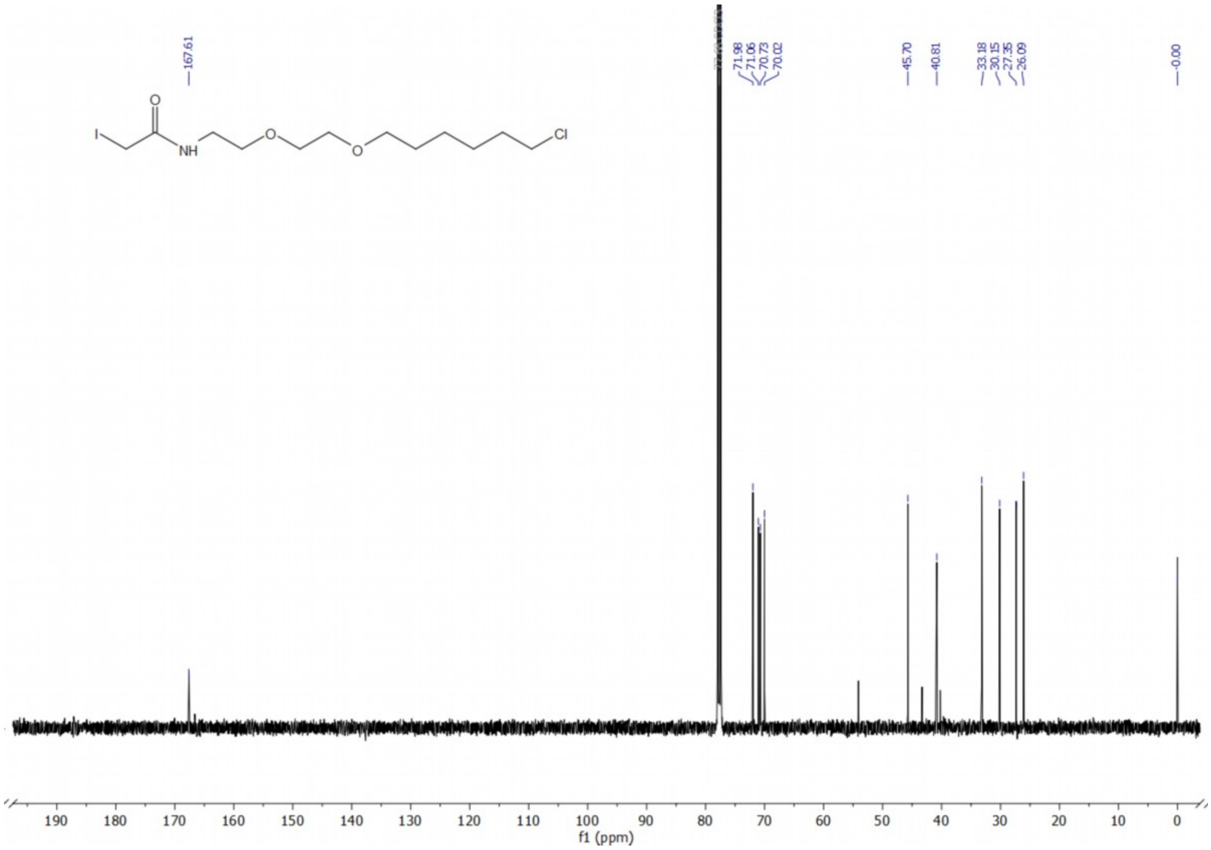

### [13]C NMR of probe 1

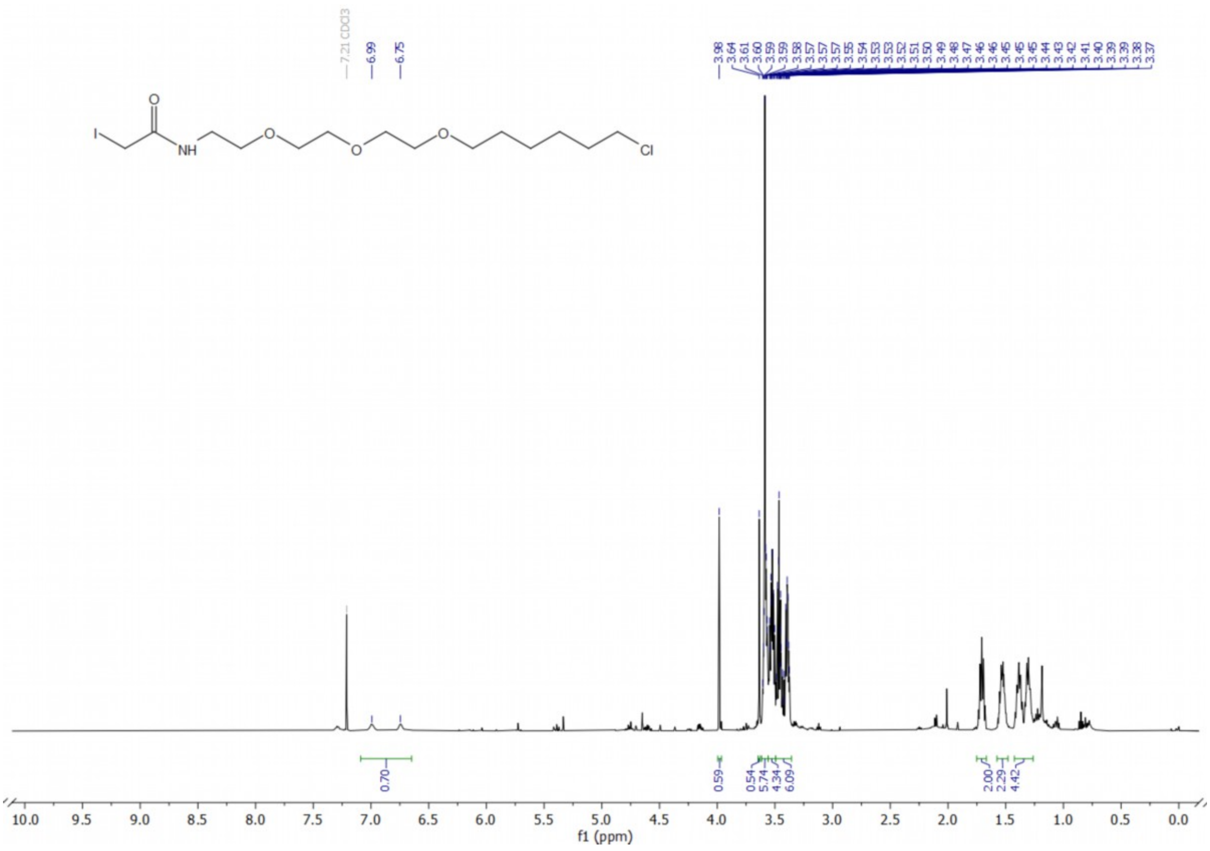

**[1]H NMR of probe 2**

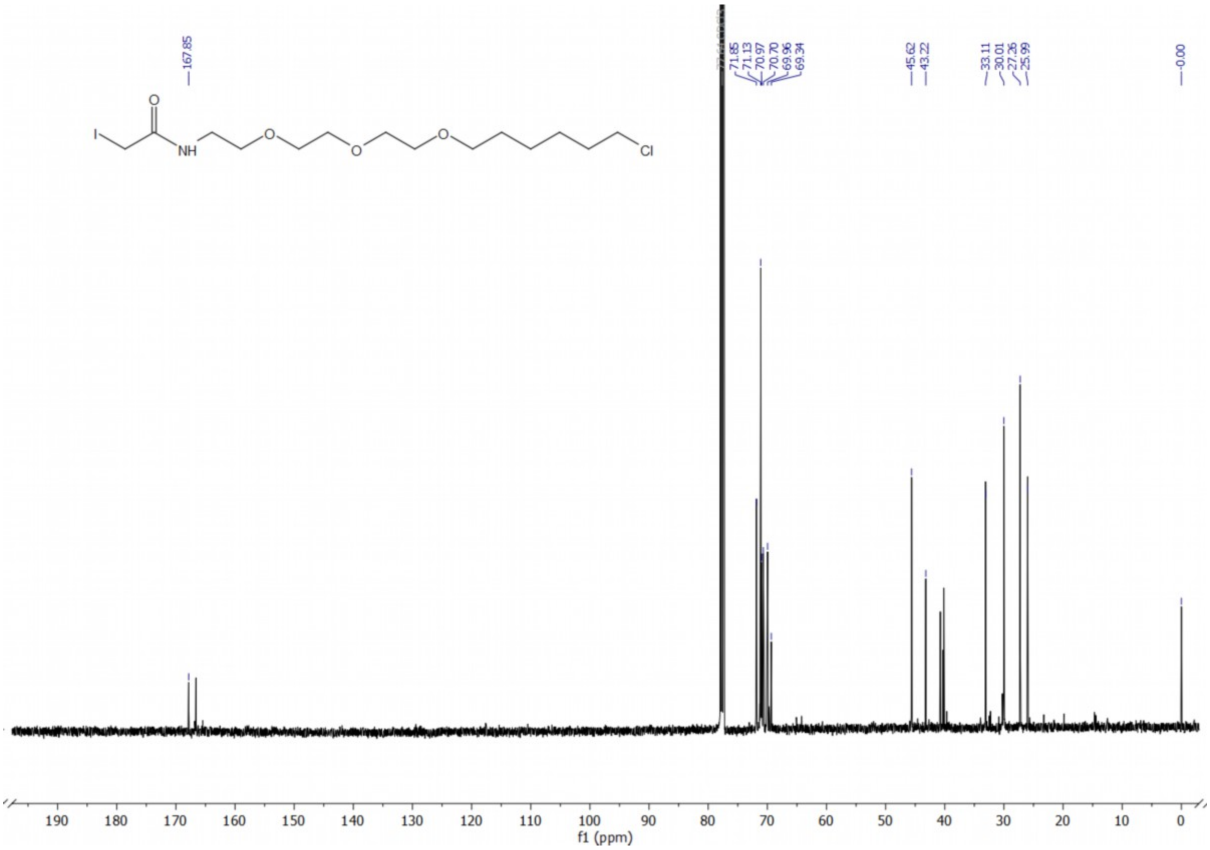

**[13]C NMR of probe 2**

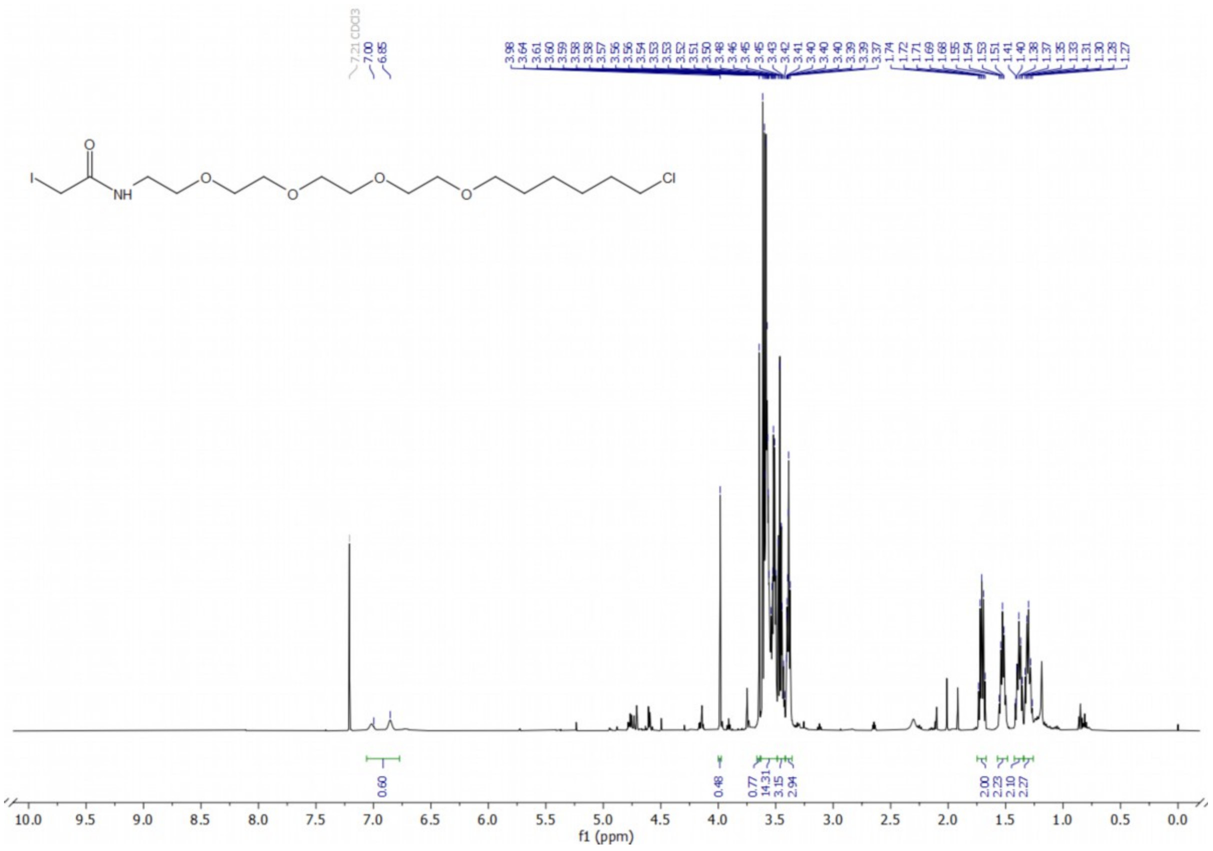

**$^{1}$H NMR of probe 3**

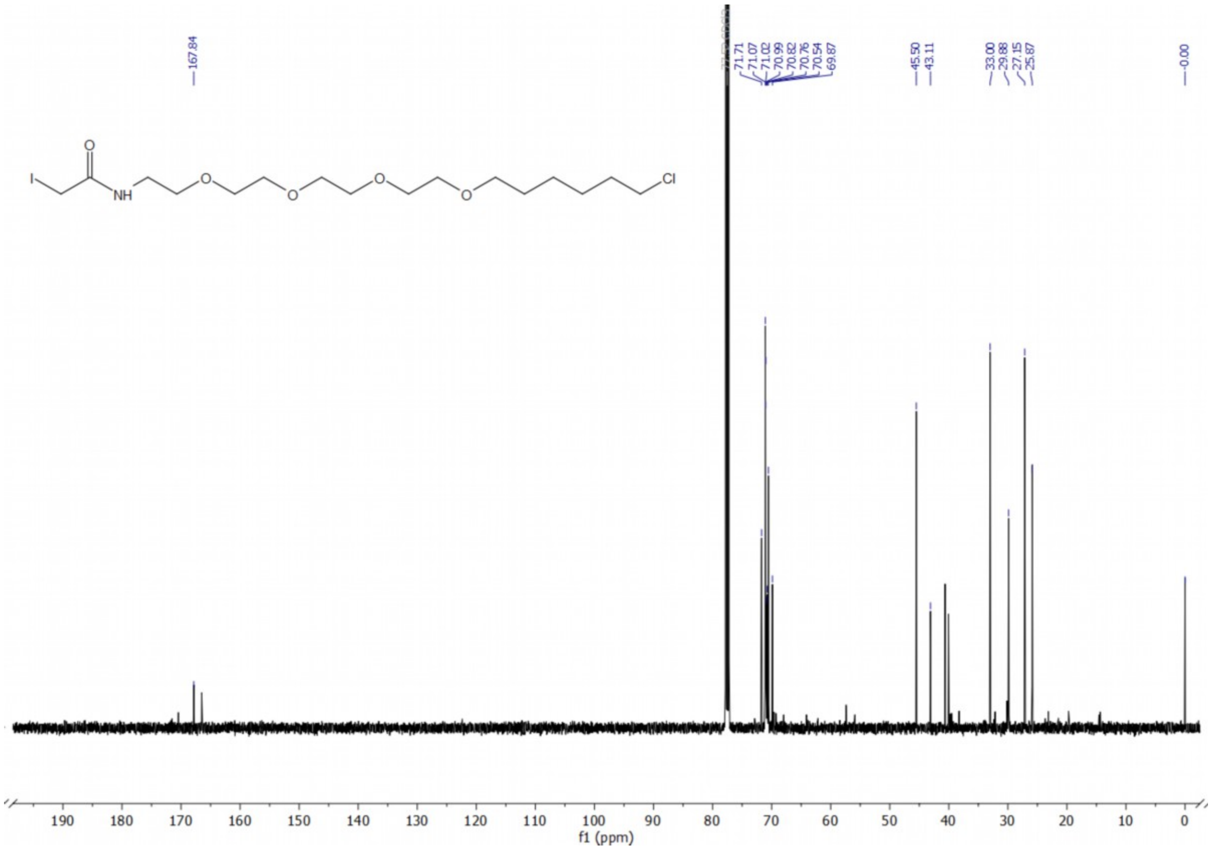

**$^{13}$C NMR of probe 3**

Uncropped Gels

**Figure 1E**

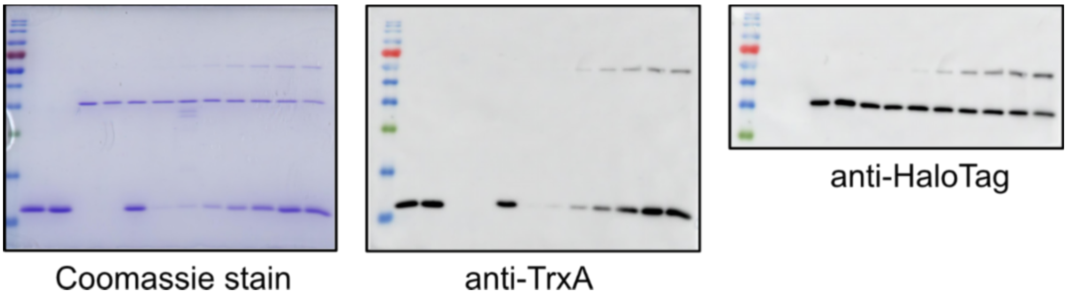

**Figure 2A**

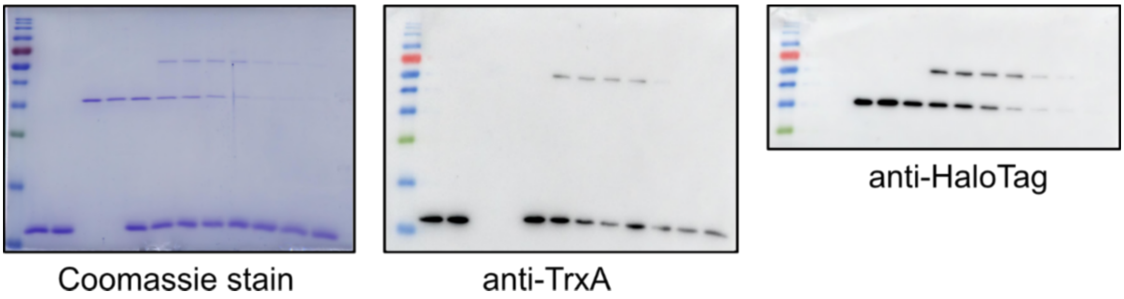

**Figure 2B**

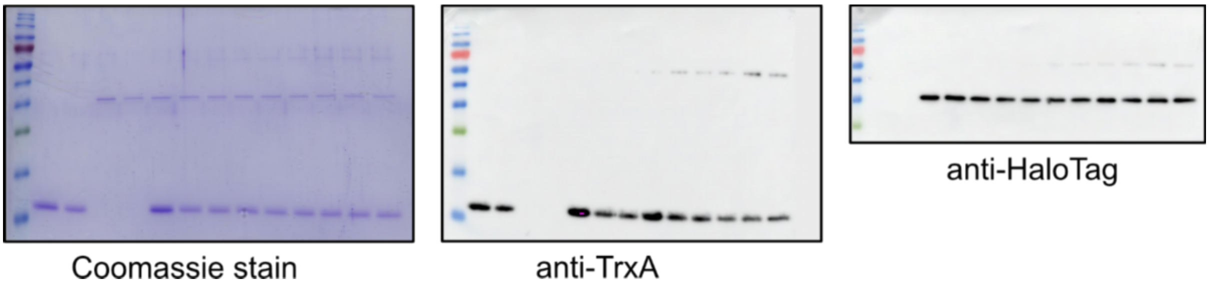

**Figure 2C**

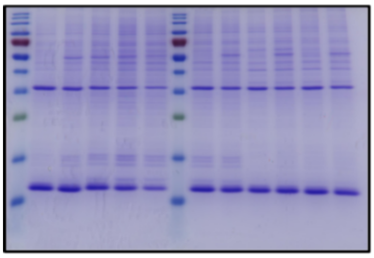

Coomassie stain

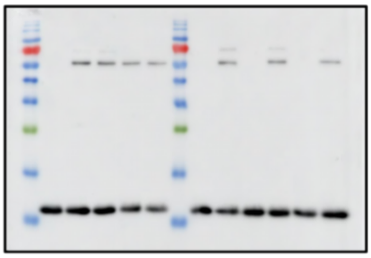

anti-TrxA

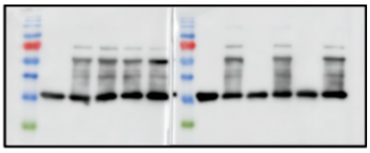

anti-HaloTag

**Figure 2D**

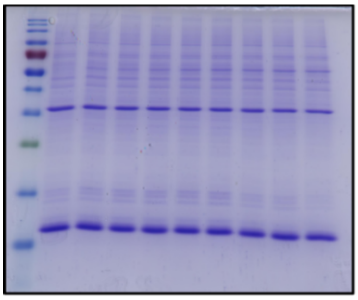

Coomassie stain

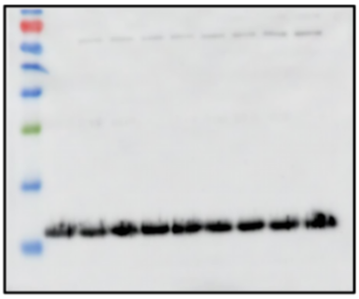

anti-TrxA

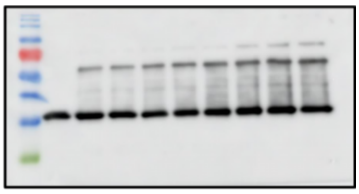

anti-HaloTag

**Figure 2E**

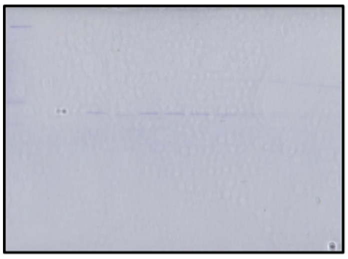

Coomassie stain

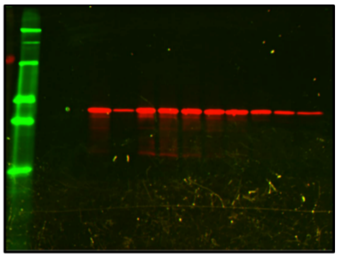

Fluorescence

**Figure 2F**

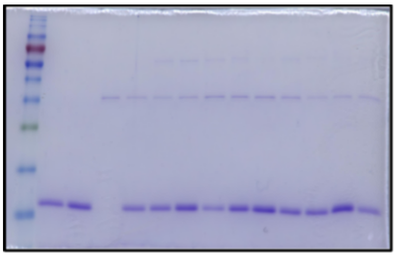

Coomassie stain

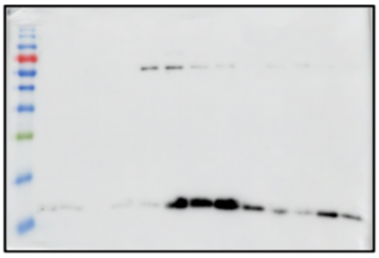

anti-TrxA

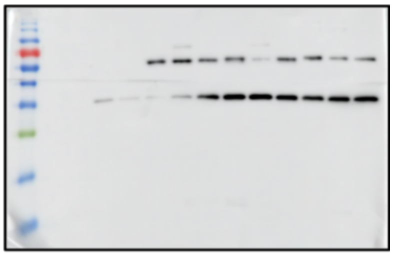

anti-HaloTag

**Figure S2**

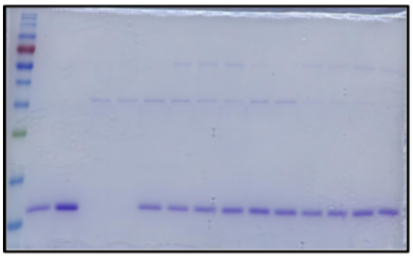

Coomassie stain

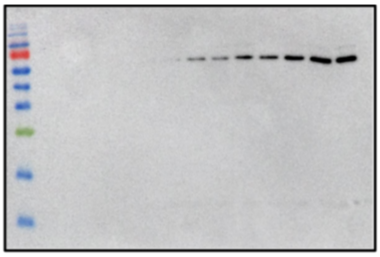

anti-TrxA

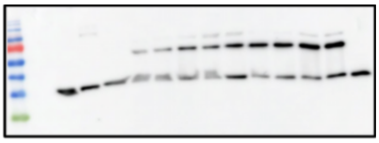

anti-HaloTag
